# MYB59 is linked to natural variation of water use associated with warmer temperatures in *Arabidopsis thaliana*

**DOI:** 10.1101/2025.02.27.640580

**Authors:** John N. Ferguson, Oliver Brendel, Ulrike Bechtold

## Abstract

Water-use contributes significantly to plant growth and productivity, yet the extent to which variation in water-use is a function of adaptive differentiation is unknown. Here, we studied natural variation of water use in *Arabidopsis thaliana* to understand how climatic history impacts water use strategies. We performed a survey of vegetative water use (VWU) and life-history traits across *A. thaliana* ecotypes in controlled and outdoor settings. Performance of select ecotypes in controlled experiments was reflective of performance in outdoor conditions. Through trait-climate and genome-wide association (GWAS) analyses, we tested for signals of environmental adaption in water use. Ecotypes from warmer environments were noted in displaying enhanced water use, independent of precipitation. GWAS identified *MYB59* as a determiner of VWU. Functionally significant *MYB59* SNPs showed associations with temperature, but not precipitation, and *myb59* mutants demonstrated reduced water use under high temperatures. Our study suggests intraspecific variation in water-use can be explained in part by climatic history, where temperature is the most significant driver. *MYB59* appears involved in this association and holds promise for study in crops.

## Introduction

Soil water availability is the most fundamental constraint to the net primary productivity of ecosystems (Craine *et al*., 2012; Zhang *et al*., 2017) and crop yields (Elliott *et al*., 2014; Lobell *et al*., 2020). This is reflected by the plethora of evolutionary adaptations that facilitate the acquisition and retention of water (Delwiche & Cooper, 2015). The trade-off between water loss and photosynthetic carbon gain is central to the constraining nature of water availability (Leakey *et al*., 2019; Brendel 2021). Thus, it follows that indices of water use efficiency (WUE) in natural systems correlate with trends in precipitation (Adams *et al*., 2019; DeLeo *et al*., 2020). Due to the lack of biochemical harnessing, the evaporation of water through stomata, i.e. transpiration, has been defined as a *loss* (Morison *et al*., 2008; Brendel 2021), despite its importance for promoting growth via numerous physiological processes (Fricke, 2017; Ferguson *et al*., 2021). In warm environments, transpiration is critical for facilitating leaf cooling, thereby stabilising photosynthesis and leaf energy budgets (Ansari & Loomis, 1959; Lambers *et al*., 1998). The degree to which transpiration or physical traits facilitate persistence in warm environments depends on water availability. However, in both hot-wet and hot-dry environments, transpiration has been demonstrated to be important for stabilising leaf temperatures across multiple species (Lin *et al*., 2017).

The natural variation present in the model species *Arabidopsis thaliana* has been well utilised to understand the physiological and genetic mechanisms that control water use and water use efficiency (WUE). For example, integrated measures of intrinsic WUE (δ^13^C; hereafter termed *W_i_*) have been employed to observe a common link with phenology (McKay *et al*., 2003, 2008; Juenger *et al*., 2005) as well as to identify genetic loci and candidate genes that regulate δ^13^C, often via stomatal physiology and behaviour (Masle *et al*., 2005; Des Marais *et al*., 2014; Campitelli *et al*., 2016). The extent to which climatic variation shapes selection on δ^13^C is complex (Brendel and Epron 2022), with findings in *A. thaliana* highlighting seasonal variation in water availability as a selective pressure on *W_i_* in non-arid environments (Dittberner *et al*., 2018). Any potential associations are difficult to dissect due to the influence of confounding effects not associated with climate, such as competition, which has been shown to play an important role in this dynamic, where lines with low WUE actually perform better under increased resource competition (Campitelli *et al*. 2016). Despite this, directional natural selection in *A. thaliana* is strongest in arid environments (Exposito-Alonso *et al*., 2019), which is reflected by the strong correlation between drought intensity and the frequency of loss of function alleles (Monroe *et al*., 2018). However, it is critical to remember that drought resistance, whether achieved through tolerance or avoidance, is not necessarily achieved by the same mechanisms as those that regulate WUE. Mechanisms that change photosynthesis or alter the overall level of stomatal conductance are little related to drought tolerance or avoidance. (Lovell et al. 2013, Kenney et al. 2014, Campitelli et al. 2016, Ferguson *et al*., 2018; Lorts and Lasky 2020; Brendel and Epron, 2022). It is therefore critical to remember that variation in *W_i_* suggests different drought response strategies, where high *W_i_* corresponds to drought sensitive, early closing stomata, whereas low *W_i_* corresponds to a drought escape strategy due to early flowering (Brendel and Epron 2022). These strategies clearly involve very different molecular or physiological mechanisms (Ferguson et al 2018), especially when biomass gain, and water loss are upscaled to the whole plant level.

Climate has been successfully determined as a selective agent across *A. thaliana* natural variation, especially along altitudinal and latitudinal clines (Stinchcombe *et al*., 2004; Wolfe & Tonsor, 2014; Vidigal *et al*., 2016; Tabas-Madrid *et al*., 2018, Sartori et al., 2019), where observed trends are often described relative to flowering time. A key exception to this is the study of (Vasseur *et al*., 2018), who observed that increasing ruderality (plants that reproduce quickly and preferentially invest resources into seed production, linked to high nutrient concentration and high net photosynthetic rate; Vasseur et al 2018, May et al 2017) was linked with mean annual temperature at sites of origin. The manner of this association highlights how ecotypes from warmer environments operate less conservatively in terms of resource use. When considering water use, this could be hypothesized to co-occur with increasing photosynthesis (Dittberner *et al*., 2018; Vasseur *et al*., 2018). Ecological shifts in *A. thaliana* have also been linked to water availability and humidity levels, suggesting that *A. thaliana* originated in dry and arid regions and adapted to high precipitation or humidity regions (Lou et al 2022). While these studies facilitate a better understanding of how general ecological strategies transect environmental gradients, they do not explicitly test how climatic selection shapes the use of any given resource, e.g., nitrogen, water, etc. It has recently been postulated that examining key traits, such as resource use, and the genes that underlie these traits in the global ecosystem context will be essential for resolving outstanding questions in ecology and molecular biology (Takou *et al*., 2019). Moreover, the applicability of such traits to crop performance may help tailor crop improvement at a regional level, since a substantial proportion of variation in breeding systems can be ascribed to climate (Ansaldi *et al*., 2018).

Present understanding of how climatic history has shaped intraspecific variation in plant water use is limited but critical for understanding determinants of community dynamics and for guiding crop development for non-benign environments. *A. thaliana* is an ideal model for this purpose due to associated germplasm and genetic resources and the wealth of climatic data from their place of origin. Arabidopsis genotypes and have been used *W_i_*, screens estimated from δ^13^C (McKay et al 2003; Juenger et al 2005; Des Marais et al 2014), which is a purely leaf level trait. However, for inferences towards breeding, it is essential to know about whole plant water use. VWU is a good proxy for that, as it encompasses whole plant traits, such as total leaf surface area, a major determinant of whole plant transpiration, which can overcompensate variations in stomatal conductance (Bogeat-Triboulot et al 2019). We have previously demonstrated the measurement/screening for water use in *A. thaliana* through short term pot drying experiments (Ferguson *et al*., 2018). The vegetative water use (VWU) measured via this approach reflects whole plant transpiration can be a useful tool for forecasting lifetime water use and we have used this to identify associated quantitative trait loci (QTL) in a biparental population (Ferguson *et al*., 2019). However, such artificial populations are not applicable for defining climate-trait interactions. To this end, we focused this study on a detailed physiological characterisation of VWU in a diverse panel of natural variants to explore how climatic history exerts selective pressures on water use in *A. thaliana* (Figure 1). In support of this, we integrated genome-wide association mapping (GWAS) with transgenic manipulation and exploration of allelic variation at a VWU candidate gene to better understand the dynamic between climatic history and water use.

**Figure 1.**
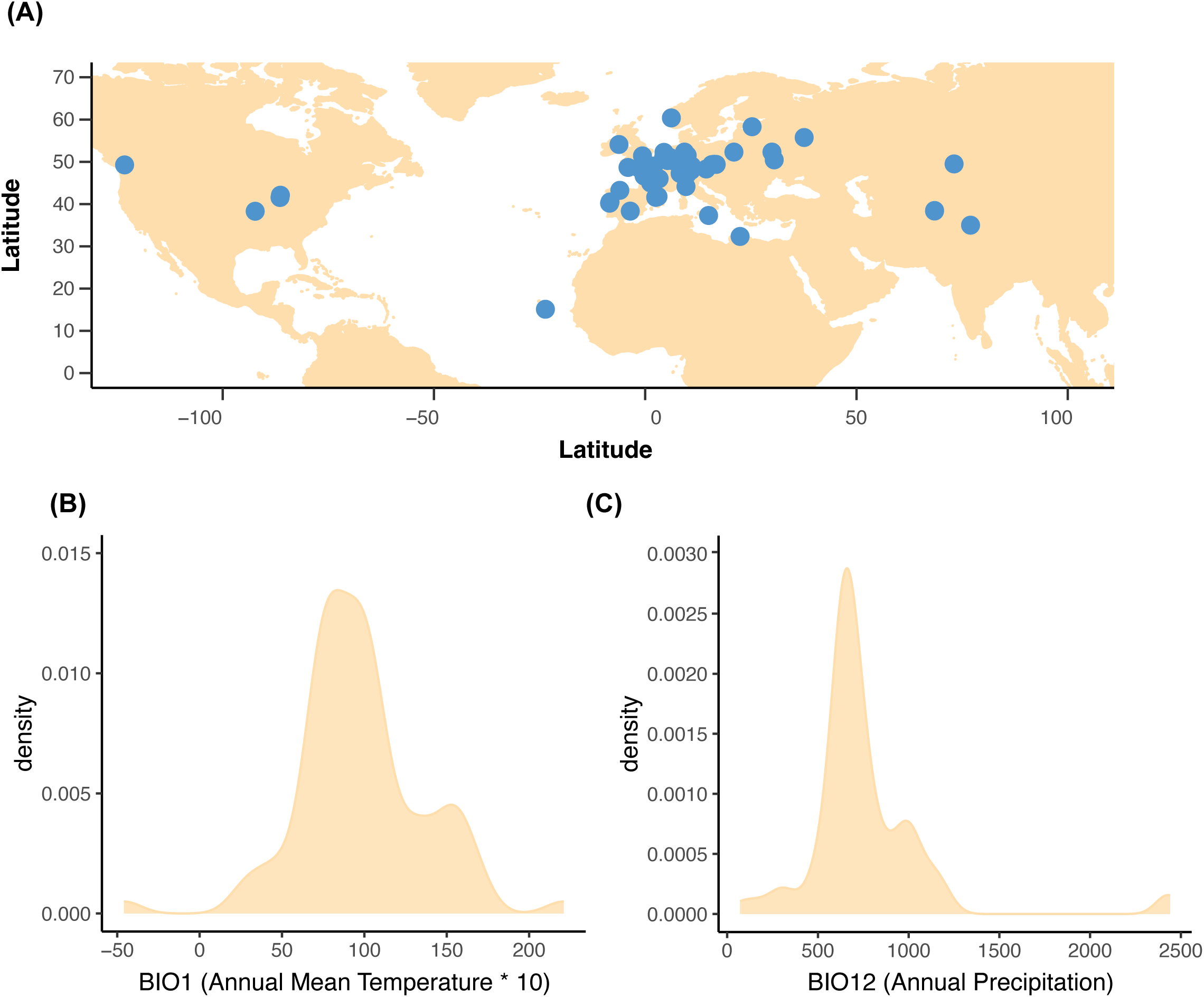
Ecotypes comprising present study (A) Geographic point of origin of all ecotypes. (B) Variation in BIOCLIM 1 (Mean annual temperature) of all ecotypes. (C) Variation in BIOCLIM 12 (Annual precipitation) of all ecotypes.

## Results

### Extensive natural variation in vegetative water use, phenology, and biomass accumulation

To study the natural variation of water-use in *A. thaliana* we subjected 78 ecotypes (Figure 1, Supplemental Table S1) to a period of water withdrawal as described previously (Ferguson *et al*. 2018, 2019). We have previously shown that this water withdrawal period does not elicit a classic ‘drought escape’ floral transitioning mechanism, nor does it reduce subsequent vegetative or reproductive biomass (Ferguson *et al*. 2018).

Although the response of relative soil water content (rSWC) to elapsed time was marginally curvilinear (e.g., Col-0 and Se-0 as shown in Figure 2A), we modelled the overall response as a linear regression (Supplemental Figure S1), and the associated slope provides an estimate of vegetative water use (VWU) over the duration of the dry down period (Figure 2B; Ferguson et al 2018). We also modelled the response of VWU to drying through a segmented approach (Figure 2C; Supplemental Figure S1), which provides an additional estimate of drought sensitivity (Ferguson et al. 2018). Importantly, the parameters estimated from this approach (initial slope (Slope 1), breakpoint, and the secondary slope (Slope 2)) were all highly correlated to VWU (Figure 2D).

**Figure 2.**
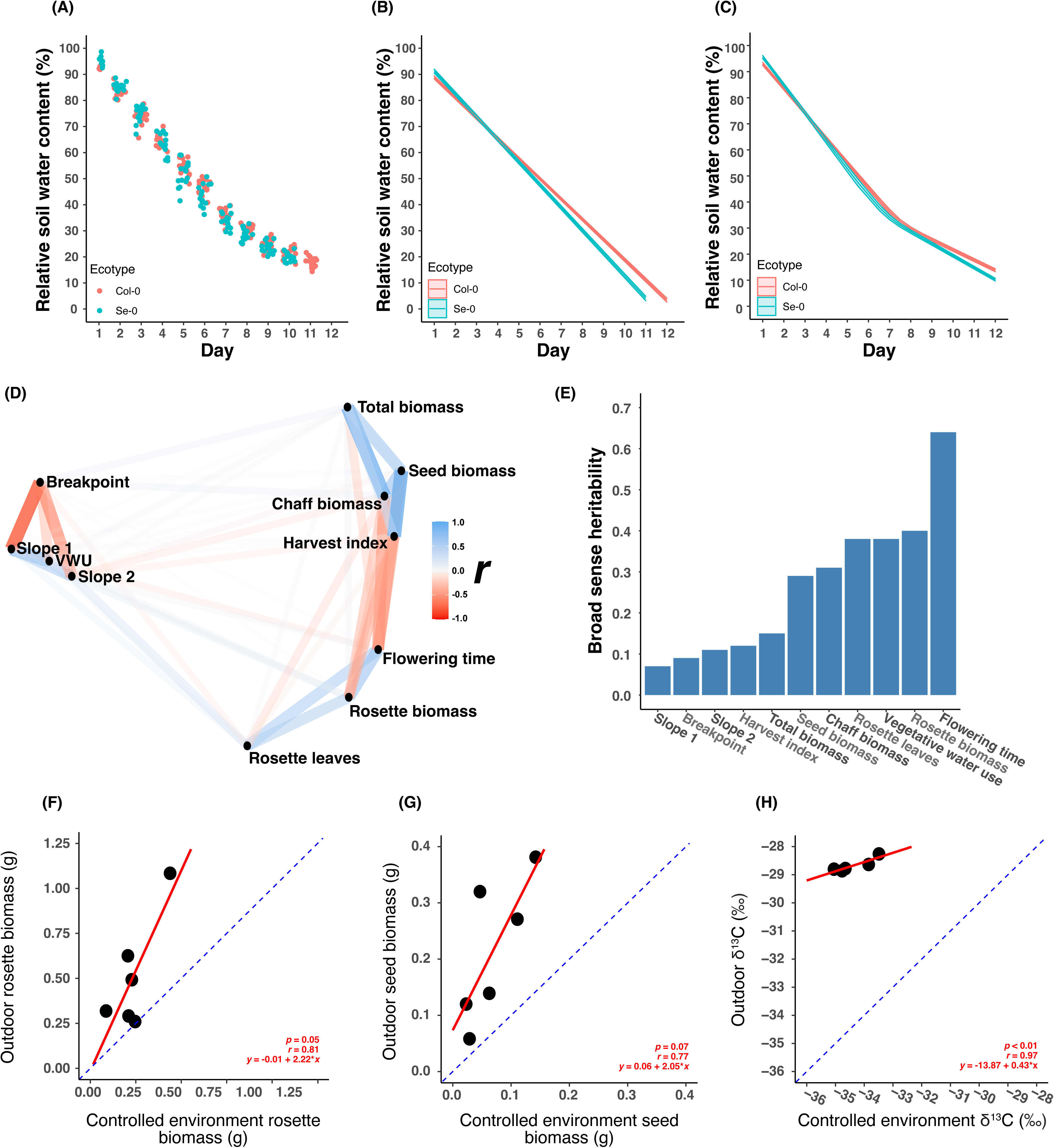
Modelling water use and trait correlations. (A) Relative soil water content (rSWC) of Col-0 and Se-0 ecotypes during the dry-down period. (B) Linear model of rSWC as a function of elapsed time in days. Vegetative water use (VWU) is defined as the slope of the linear model. (C) Segmented model of rSWC as function of elapsed time in days. (D) Network plot showing the relationship between all traits measured as part of the controlled experiment. (E) Broad sense heritabilities (*H^2^*) of all traits measured. (F) Correlation between vegetative biomass measured in the controlled and outdoor experiments. (G) Correlation between reproductive biomass measured in the controlled and outdoor experiments. (H) Correlation between carbon isotope discrimination measured in the controlled and outdoor experiments.

VWU demonstrated a 2.5-fold variation across the surveyed diversity (Supplemental Figure S2; Supplemental Table S2) and moderate *H^2^* (Figure 2E), suggesting a discernible genetic component to the observed variation.

Across the diversity surveyed, we observed substantial variation in phenology (Supplemental Figure S2; Supplemental Table S2). Flowering time ranged from 45 to 105 days (Supplemental Figure S2; Supplemental Table S2) and demonstrated the highest *H^2^* of all traits comprising this study (Figure 2E). Life-history traits such as rosette biomass, seed biomass, harvest index and total above ground biomass also showed substantial natural variation (Supplemental Figure S2; Supplemental Table S2) and demonstrated low-to-moderate *H^2^* (Figure 2E).

Using data obtained from our controlled environment study (Supplemental Table S2), we explored associations between all measured traits via concurrent multidimensional clustering and Pearson product moment correlation tests (Figure 2D; Supplemental Table S3). Traits related to water use, biomass accumulation, and phenology were observed to independently cluster together (Figure 2D). Traits relating to water use, i.e., VWU, Slope 1, and Slope 2, were positively correlated with each other and negatively correlated with the breakpoint estimated via the segmented modelling approach (Figure 1D; Supplemental Table 3). VWU did not demonstrate a significant association with any further traits (Figure 2D; Supplemental Table 3). This suggests that short-term water use was not driven by development or eventual biomass accumulation. However, across a separate experiment comprising 48 ecotypes in the same controlled conditions, we did observe a significant positive association between rosette area (measured prior to the drying period used to estimate VWU) and VWU (Supplemental Figure S3) which is line with previous findings (de Ollas et al. 2023).

We observed a trade-off between time to reproduction and reproductive output (Figure 2D; Supplemental Table S3). Flowering time was strongly, negatively correlated with the mass of chaff (stalks and pods) and seed biomass (Figure 2D; Supplemental Table S3). This suggests that in this environment, ecotypes that delay floral transitioning have reduced fitness. Conversely, later flowering ecotypes tended to have much larger vegetative rosettes, highlighting a floral-mediated trade-off between vegetative and reproductive growth (Figure 2D; Supplemental Table S3).

### Comparison of life-history traits measured in controlled growth room versus common garden environments

A major aim of this study was to test for phenotype-climate associations, we assessed whether plant performance in our controlled growth room experiments was reflective of performance in a *more natural* outdoor experiment (Supplemental Table S4). For this, we grew a set of six ecotypes (C24, Col-0, Ct-1, Est-1, HR-5, and Se-0) in both environments.

The performance of these ecotypes was determined via the accumulation of biomass (above ground vegetative and reproductive biomass) and via integrated water use efficiency (measured as leaf carbon isotope discrimination (δ^13^C)). In all three cases, the performance of ecotypes in controlled and outdoor environments were positively correlated (Figure 2F-H), highlighting that life-history and water use traits measured in controlled environments can be reflective of those some traits phenotyped in more natural settings.

Taken together, the findings from these comparative analyses suggests that phenotypes measured in controlled environments can be representative of those measured in more dynamic and variable growth conditions, such that it is feasible to identify links between climatic parameters at the point of origin of ecotypes (Supplemental Table S1) and phenotypic traits assessed in controlled environments (should such associations exist). Additionally, these results demonstrated that the outdoor environment was favourable for the ecotypes tested here, since vegetative and reproductive biomass was overall enhanced (Figure 2F-G).

### Associations between climatic variation and phenotypic variation

To test for trait-climate associations we adopted the comparative principal component analyses (PCA) approach utilised by Wolfe and Tonsor (2016), where trait principal components (TPCs) are compared to climate principal components (CPCs). It allows to extract critical information from a complex set of phenotypic traits by reducing a larger set of initially correlated variables to a much smaller set of uncorrelated variables, while preserving information.

The trait PCA (Figure 3) confirmed previous observations regarding the associations of water use, phenology and biomass accumulation associated traits (Figure 2D; Supplemental Table S3). The first three TPCs were observed to explain the majority of the observed variation (73.7%; Supplemental Figure S4A). TPC1 explained 33.4% of the total trait variation and was predominantly associated with phenology (negative correlation) and biomass accumulation (positive correlation for reproductive and total biomass and negative correlation for rosette biomass) (Figure 3C). TPC2 explained 25.2% of the total trait variation and was predominantly associated with water use (positive correlation) and drought sensitivity (i.e., breakpoint, negative correlation) (Figure 3C). TPC3 was characterised by positive associations with all biomass accumulation traits (Figure 3C).

**Figure 3.**
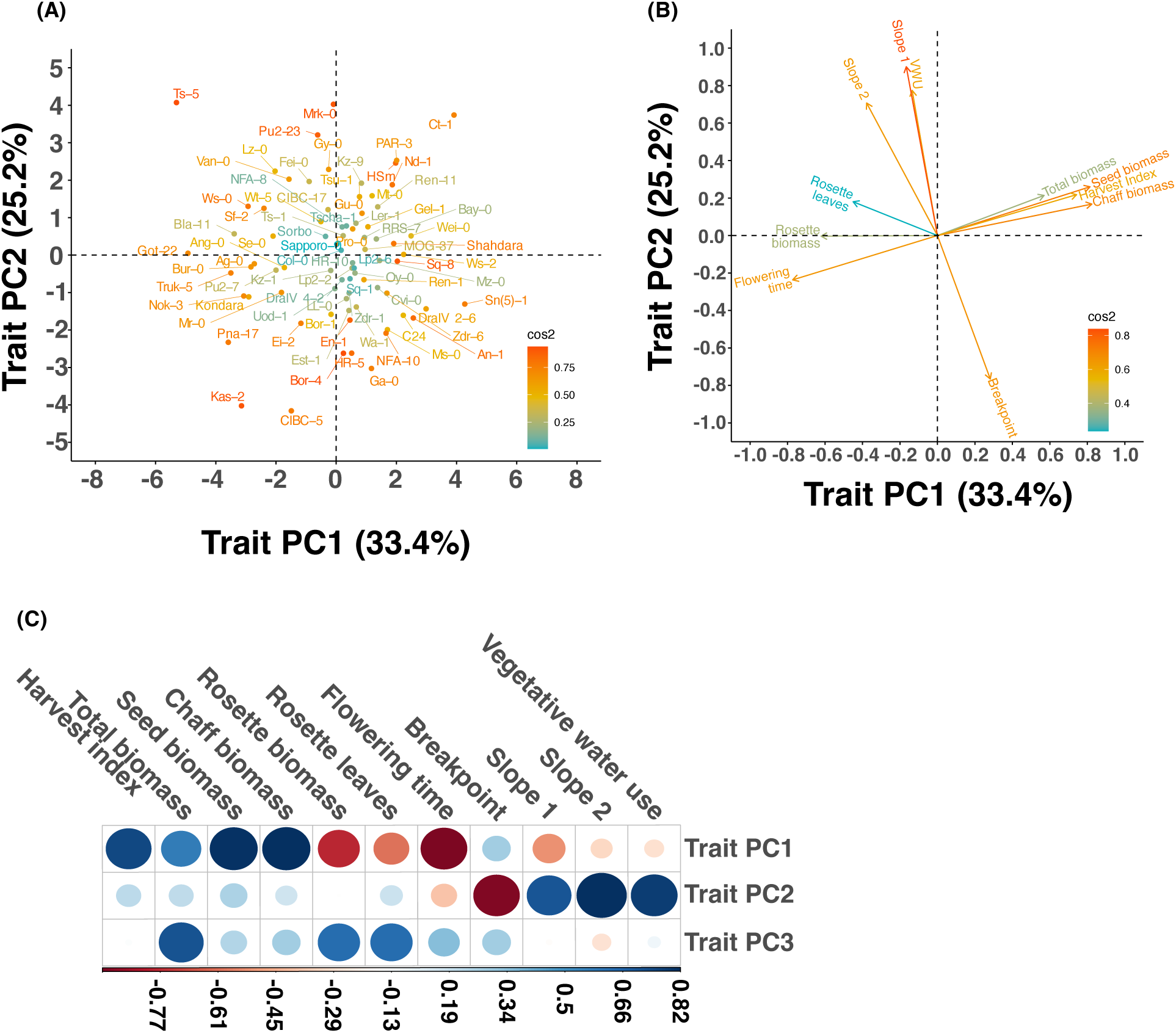
Trait principal component analysis (A) The loading of all traits onto TPC1 and TPC2. Traits are coloured according to the quality of the factor map (cos^2^). (B) The association between all ecotypes with TPC1 and TPC2. Ecotypes are coloured according to cos^2^. (C) Relationship between all traits with the first three TPCs. The size of each bubble is proportional to the strength of the association and the colour represents the direction of the association as indicated by the colour bar.

For the climate PCA (Figure 4), we observed that the parameters related to precipitation, temperature, and seasonality tended to cluster with one another, with a few exceptions (e.g., BIO8 and BIO2; Figure 4A). The first five CPCs explained the majority of the total climatic variation (90.1%, Supplemental Figure S4B). CPC1 explained 32% of the total variance and was most strongly associated with temperature (positive correlation) and temperature seasonality (negative correlation) (Figures 4A-C). A key for the BIOCLIM parameters is shown in Figure 4D.

**Figure 4.**
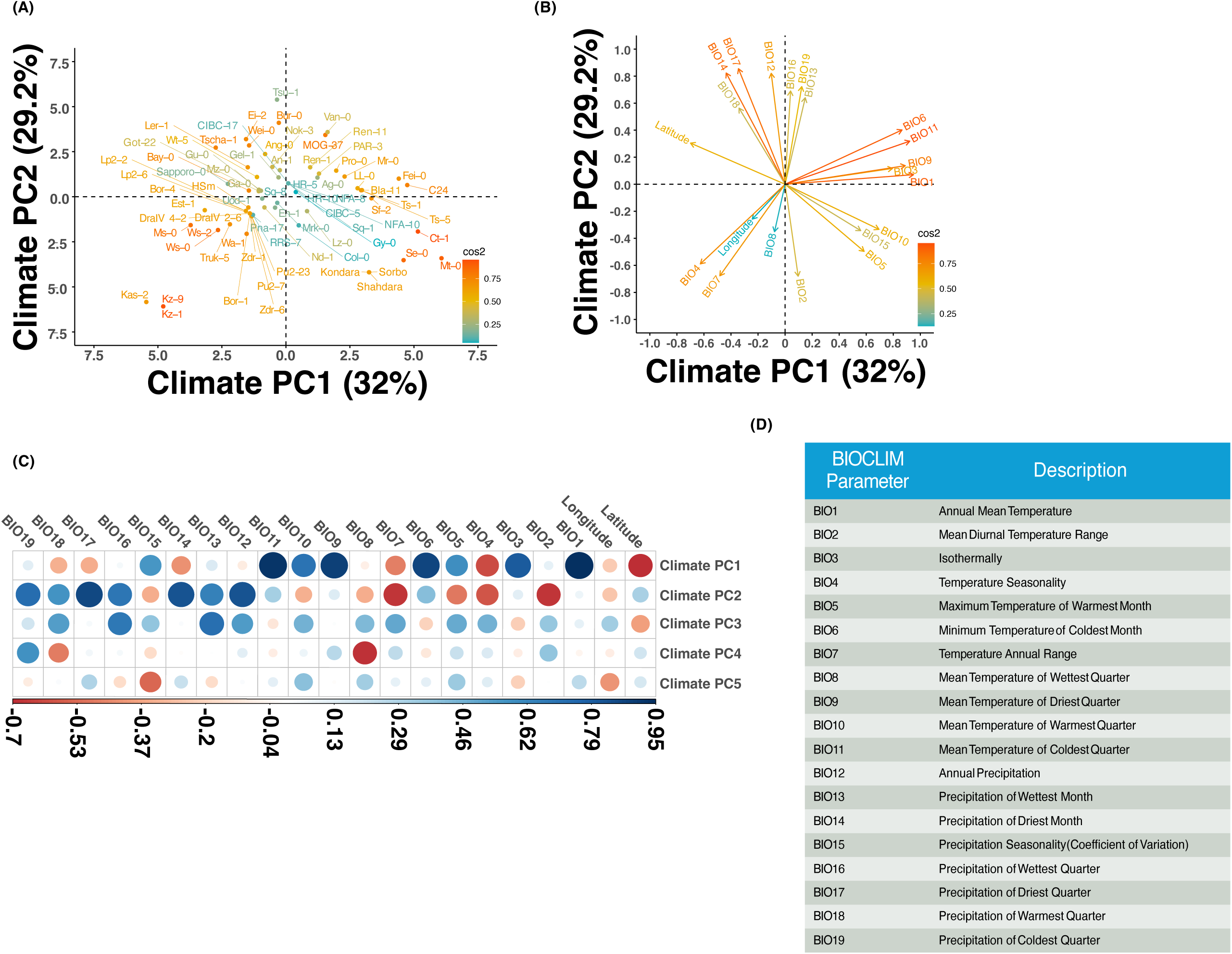
Climate principal component analysis. (A) The loading of all traits onto CPC1 and CPC2. Climate parameters are coloured according to the quality of the factor map (cos2). (B) The association between all ecotypes with CPC1 and CPC2. Ecotypes are coloured according to cos2 (C) Relationship between all traits with the first five CPCs. The size of each bubble is proportional to the strength of the association and the colour represents the direction of the association as indicated by the colour bar. (D) Definition of BIOCLIM parameters used in the CPCA.

To look for signals of climatic selection on plant performance, we subsequently tested for associations between the significant TPCs and CPCs (Supplemental Table S4) following the approach of Wolfe & Tonsor (2016). Of the 15 possible association tested here, we only observed a significant association between CPC1 and TPC2, where the association was positive (Figure 5A, Supplemental Table S5). This suggests that ecotypes with increased VWU originate from environments that are consistently warmer throughout the year (i.e., warm environments with reduced seasonality).

**Figure 5.**
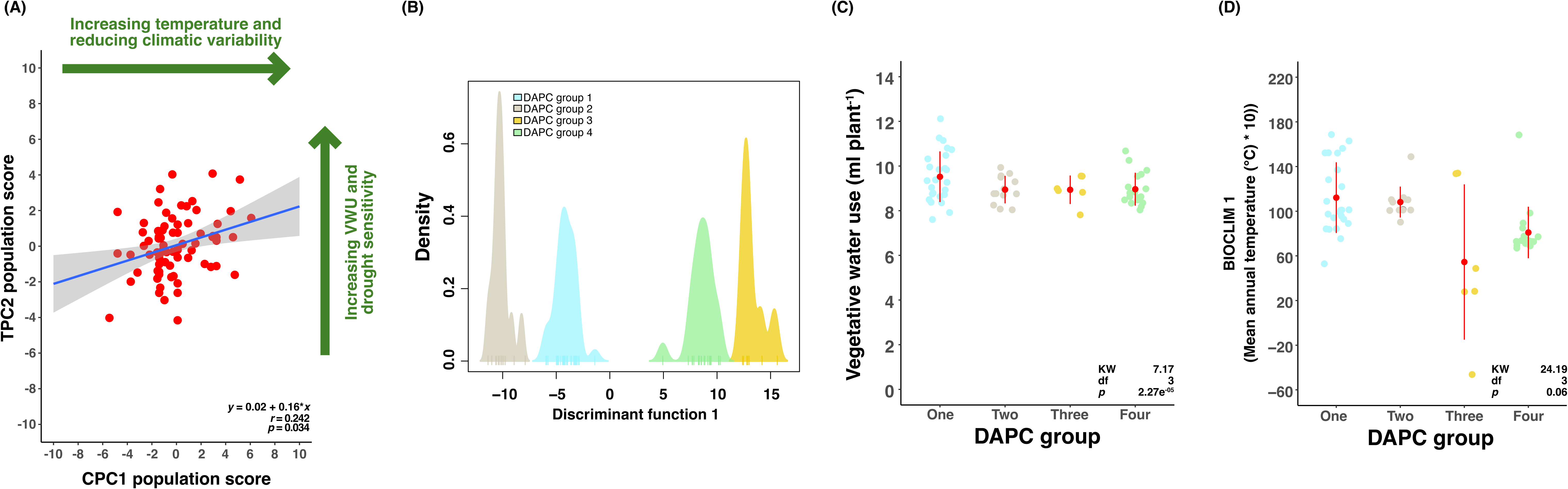
Association between temperature and water use. (A) Correlation between the ecotype population score for climate principal component 1 (CPC1 population score) and the ecotype population score for trait principal component 2 (TPC2 population score). The linear regression of the relationship (blue) and standard error of the model (grey) are plotted. The Pearson correlation coefficient (*r*) and associated p-value cut-off are inset. The most important parameters associated with CPC1 and TPC2 are highlighted in green to aid interpretation. (B) Summary of discrimination analysis of principal components (DAPC) to group ecotypes based on genetic similarity. Each tick on the discrimination function 1 axis represents an individual ecotype. Each cluster of ecotypes is named and coloured uniquely. (C-D) Vegetative water use and mean annual temperature of all ecotypes grouped according to their DAPC classification. Kruskal-Wallis test statistics are inset. Red points and error bars denote the DAPC group mean and standard deviation.

We additionally clustered the ecotypes according to genetic similarity via discrimination analyses (Figure 5B). Comparing VWU and BIOCLIM 1 (mean annual temperature) across the distinct genetic ecotype clusters demonstrated that both parameters were significantly different across these groups (Figure 5C-D). Moreover, the two groups that clustered negatively (DAPC group one and two) onto the first discrimination function (which explains the majority of the genetic variation) demonstrated higher VWU (Figure 5C) and were from warmer environments (Figure 5D) compared to the two groups that clustered positively (DAPC group three and four) onto this first discrimination function. This suggests that there may be an adaptive genetic basis to the above-described association between temperature and water-use.

### Identification of *MYB59* as a candidate gene for vegetative water use

We performed GWAS to identify SNPs significantly associated with the variation observed for VWU (Figure 6A). The AMM GWAS approach identified two significant SNPs in close proximity to one another. These SNPs were at base-pair position 24088050 (-log_10_(*p-value*) = 6.77) and 24088195 (-log_10_(*p-value*) = 6.43) on chromosome five (Figure 6A, Supplemental Table S6). Within the LD block (R^2^ > 0.3) containing the two significant SNPs there was just a single gene, transcription factor *MYB59* (AT5G59780; base-pair coordinates: 24081922-24083562; Supplemental Figure S5).

**Figure 6.**
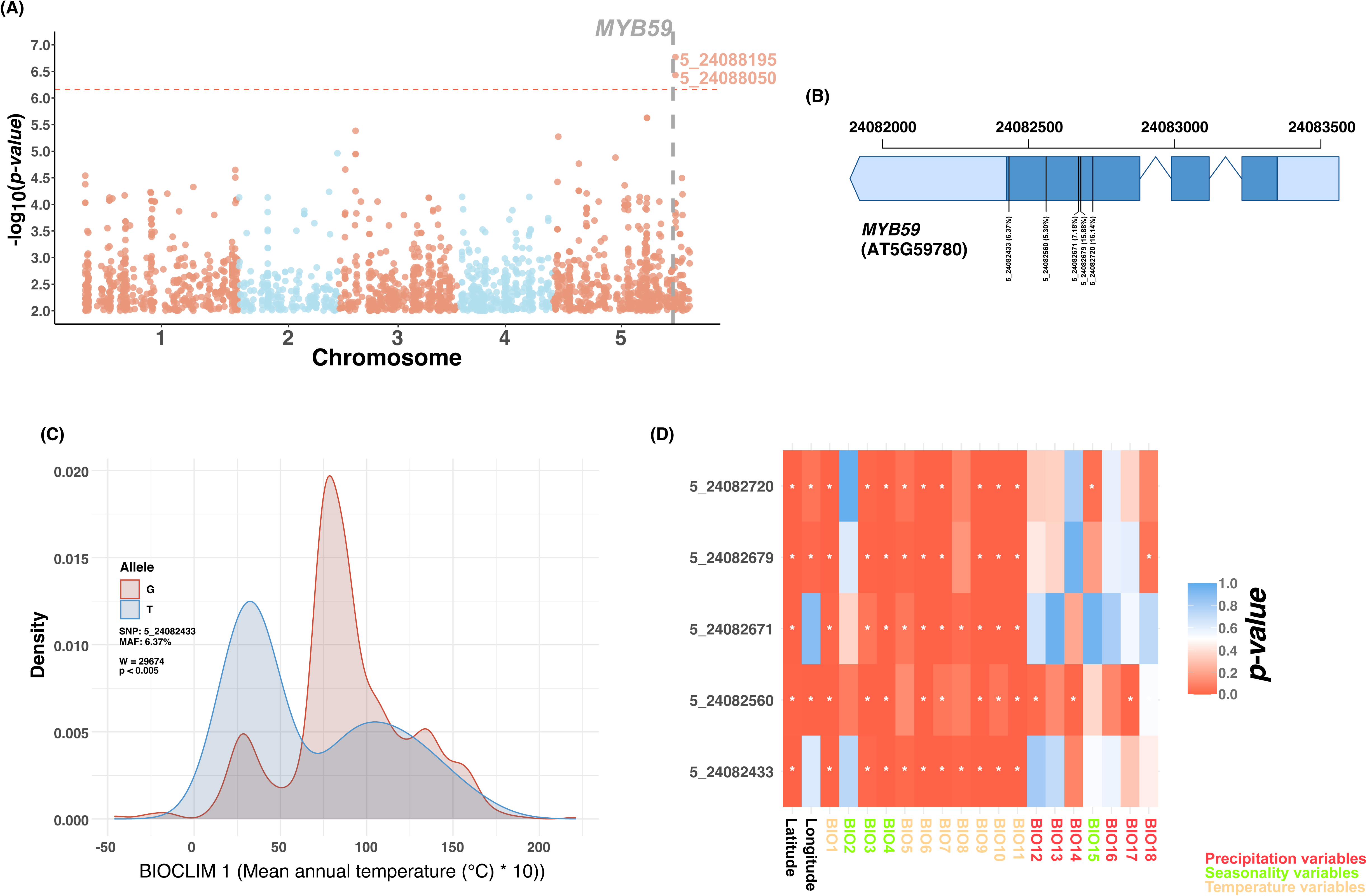
GWAS for vegetative water use (VWU) and exploration of allelic variation at *MYB59*. (A) Manhattan plot showing the results of GWAS for VWU. A suggestive cut-off 6 -log_10_(*p-value*) is highlighted, where the two SNPs above this threshold have FDR below 0.05. The position of *MYB59* is indicated. (B) *MYB59* gene model with position on chromosome 5 indicated. Light blue regions indicate untranslated regions. Dark blue regions indicate exons. Thin blue lines indicate introns. The location of SNPs with predicted modifying effects on the protein structure are highlighted. (C) Probability density of the two alleles of SNP 5_24082433 against BIOCLIM parameter 1. The statistics of a two-paired Wilcoxon test are inset. (D) Heat map showing the *p-values* from two-paired Wilcoxon tests where allelic variation at each of the *MYB59* functional SNPs is tested against latitude, longitude and the 18 BIOCLIM parameters. The BIOCLIM parameters are coloured according to whether they represent temperature, precipitation, or seasonality variables.

The two major haplotypes of *MYB59* (Supplemental Table S7) showed significant differences in VWU (Supplemental Figure S6A), supporting our assertion of *MYB59* as a candidate gene for regulating VWU. Furthermore, these haplotype groups did not significantly differ in rosette biomass (Supplemental Figure S6B) or flowering time (Supplemental Figure S6C), thereby suggesting that the variance in VWU imparted by *MYB59* was not linked to any impact of *MYB59* on phenology or overall plant size. The only other parameter that demonstrated a difference was Slope 1, which represents the initial portion of the dry down period (Supplemental Table S7). Additionally, we observed that the mean annual temperature (BIOCLIM 1), maximum temperature of the warmest months (BIOCLIM5) and mean temperature of the warmest quarter (BIOCLIM10) at the point of ecotype origin was significantly different between the two haplotype groups (Supplemental Figure S6D; Supplemental Table S7). Neither mean annual precipitation (BIOCLIM 12) nor any other precipitation parameters (BIOCLIM 13-18) were significantly different between the two main haplotype groups (Supplemental Figure S6E, Supplemental Table S7). This highlights how temperature, but not precipitation, may exert a selective pressure on the allelic form of *MYB59*.

To test the link between climate and *MYB59* variation we queried the Polymorph 1001 genomes database for SNPs with moderate-to-high modifying impacts on the translated amino acid sequence of *MYB59*. Through this approach we identified multiple SNPs, all of which were defined as having moderate modifying impacts on the amino acid sequence. Five of these SNPs had a minor allele frequency (MAF) greater than 5%, which ranged from 5.30%-16.14% (Figure 6B). We tested for associations between allelic frequency of these SNPs and climate at ecotype point of origin. The missense variant of the SNP at base-pair position 24082433, for example, was observed to cluster in cooler environments (Figure 6C). In general, when comparing allelic frequency of all five SNPs against all 18 BIOCLIM parameters and latitude and longitude, we noted that significant differences in the probability densities of associated alleles were restricted to temperature variables, including seasonality (Figure 6D), suggesting that temperature, and not precipitation or climate variability, exerts selective pressure on the allelic form of *MYB59*.

### *MYB59* affects water use under elevated temperature conditions by altering root length and stomatal densities

To test the importance of *MYB59* for regulating water use in warm environments we utilised two independent *myb59* knockout lines alongside wildtype. Here, we phenotyped VWU under control (22°C) and warm (27°C) conditions. In general, VWU increased under the warm conditions relative to the control treatment (Figure. 7A). However, VWU was significantly reduced in the *myb59* knockout lines relative to the wildtype under the warm conditions, but not the control conditions (Figure 7A). These differences could have been due to the marginally but not significantly increase in rosette areas between Col-0 and both *myb59* alleles (Supplemental Figure S7). We also observed that the *myb59* knockout lines demonstrated significantly reduced leaf water potential under warm conditions (Figure 7B), which is indicative of perturbed water use and water status in these lines relative to wildtype.

**Figure 7.**
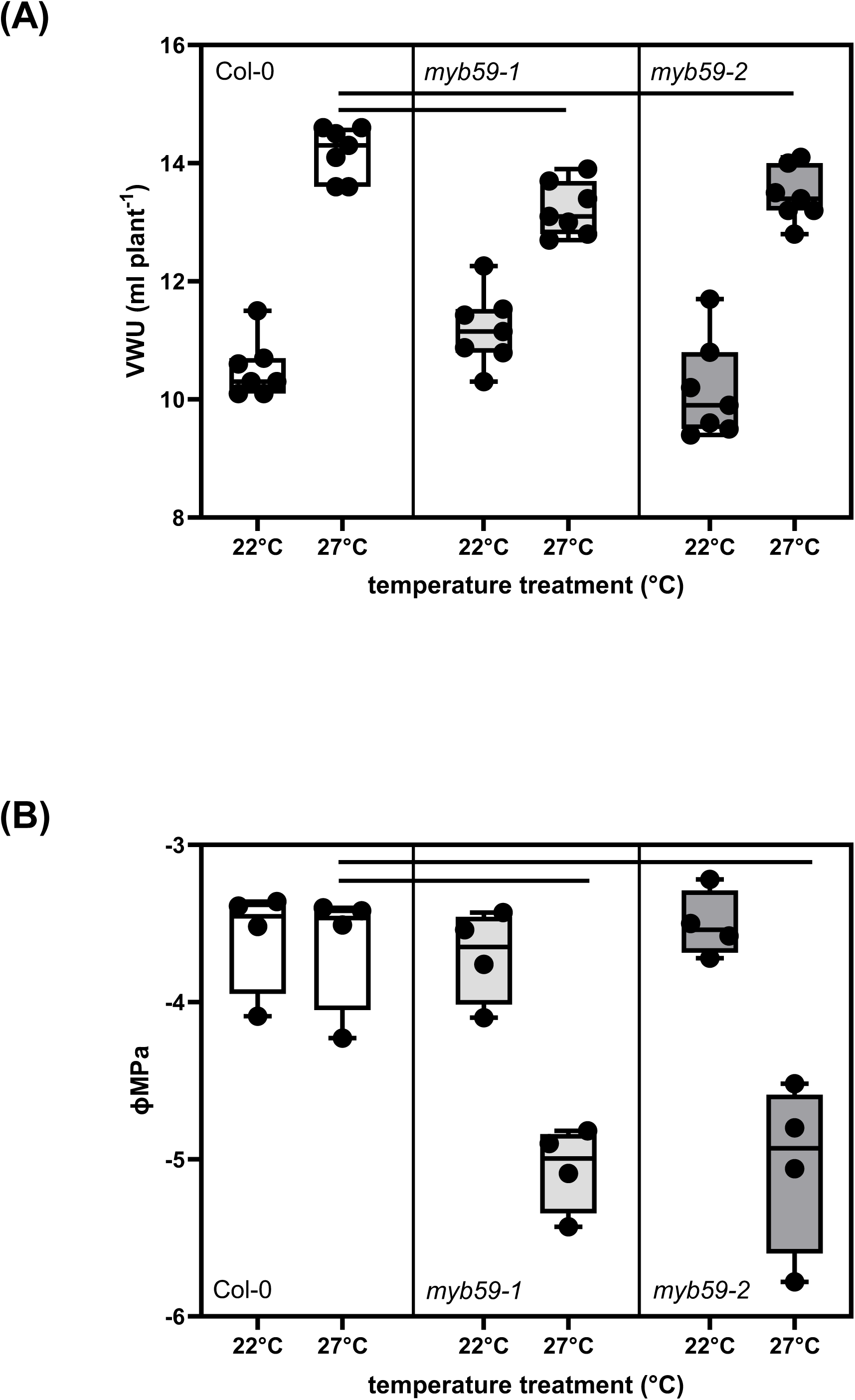
The role of *MYB59* on water use-associated traits. (A) Vegetative water use (VWU) of wildtype and *myb59* knockout lines under control and warm temperatures. (B) Leaf water potential of wildtype and *myb59* knockout lines under control and warm temperatures. Horizontal lines indicate significant difference compared to Col-0 (A) Col-0 vs *myb59-1* P < 0.0025; Col-0 vs *myb59-2* P= 0.019 and (B) Col-0 vs *myb59-1* P= 0.0025; Col-0 vs *myb59-2* P = 0.0027 (One-Way ANOVA, posthoc Tukey)

Importantly, stomatal density, which is known to be influenced by *myb59* (Fasani *et al*. 2019), showed a significant reduction at both temperature treatment (Supplemental Figure S8), yet VWU was only reduced under warm temperatures (Figure 7A), suggesting that differences in stomatal patterning and/or behaviour are unlikely to underly the observed differences in VWU under warm temperatures. Often, a change in stomatal density is accompanied by a change in cell size, which may not change the overall transpiration capacity.

Since water use is also a function of root architecture and *MYB59* is known to influence root traits (Du *et al*. 2019), we additionally profiled root length of the two *myb59* knockouts and wildtype under the elevated temperature treatment. Root length in the *myb59* knockout lines was increased when the plants were growing under the warm treatment (Supplemental Figure S9). The difference in root length between the *myb59* knockout lines and wildtype was further reflected by differences in root biomass and the root:shoot ratio (Supplemental Figure S10).

## Discussion

Classical evidence from crop species has highlighted how artificial selection in warm environments has inadvertently selected for heat avoidance via transpiration (Radin et al 1994; Koester et al., 2016). Despite this, little evidence exists to determine which selective agents define intraspecific variation of water use in natural populations. In this study, we explored natural variation of VWU in *A. thaliana* and tested for associations between this variation and climatic variation at the site of origin of the ecotypes comprising the study. The variation in VWU observed was heritable (Figure 1E) and the response of relative soil water content during the short-dehydration experiments appears coordinated to known transcriptomic responses with respect to the breakpoint in the relationship between rSWC and time and known changes in gene expression during the same experimental dry down period (Bechtold *et al*. 2016). Taken together, these findings highlight a strong genetic component to VWU variation in *A. thaliana*. This dataset builds upon a large body of work that describes natural variation of traits that underpin water use in *A. thaliana* (Dittberner *et al*., 2018; Ogura *et al*., 2019; Bhaskara Badiger *et al*., 2022; Elfarargi *et al*., 2023), by measuring water use across an extended vegetative period.

Across the ecotypes comprising this study, we observed the classical trade-off between investment in vegetative biomass accumulation and reproductive output (Figure 1D) that is well-understood to be mediated by flowering time (Ågren *et al*., 2013; Ellis *et al*., 2021)). Biomass accumulation associated to the vegetative rosette is a good approximation of plant size (Ferguson *et al*. 2018) and we observed significant variation across the surveyed ecotypes to this end (Figure 1E; Supplemental Figure S2). Plant size is often associated with enhanced water use (Feldman *et al*., 2018), however in our study we did not observe a significant association between rosette biomass and VWU or any of the other water use-associated traits (Figure 1D). This is an important observation for this study, as it suggests that VWU variation is not a function of size. Therefore, any genetic or climatic association with VWU are unlikely to be a function of variation in flowering time. This is important because flowering time is under strong selection to adapt populations to local climate (Beckerman *et al*., 2002; Ågren *et al*., 2017) and is therefore often associated with climatic variation (Stinchcombe *et al*., 2005). Consequently, the independence of VWU from phenological variation here guards against the identification of spurious association between VWU and climate parameters or SNP markers that may otherwise have had arisen had VWU been linked to biomass accumulation.

Most of our understanding on plant growth and development in *A. thaliana* has come from controlled environments. Here, we have shown that performance of ecotypes growing in our controlled environment studies matches up with performance from more natural settings (Figure 1F-H). These rank correlations between controlled and outdoor environments are supportive of exploring associations between phenotypic variation assayed in controlled environments and climatic variation at the point of origin of ecotypes. Previous work in this area has produced evidence highlighting climatic selection on traits that underpin growth and development. Through comparing TPCs and CPCs Wolfe and Tonsor (2015) demonstrated selection for increased investment in vegetative growth with reduced temperature and increased precipitation across 48 ecotypes that span a relatively small altitudinal gradient in north-eastern Spain. Similarly, Dittberner *et al*. (2018) directly compared traits with environmental PCs to show associations between WUE and stomatal characteristics, most notably they demonstrated a positive association between stomatal size and temperature. Through comparing climate-trait principal components and corroborating those results with differences in climate and trait parameters between genetic clusters of ecotypes, our study suggests that variation in VWU is in part a result of selective pressure associated with temperature (Figure 4). This finding is in accordance with reciprocal transfer experiments that have demonstrated a developmental switch to enhanced water use to promote cooling capacity in *A. thaliana* (Crawford et al., 2012).

*A. thaliana* is particularly sensitive to temperature changes (Ågren et al., 2012; Clauw et al 2022). Due to its influence on growth, temperature has been demonstrated to be key in determining the distribution of *A. thaliana* (Agren *et al* 2012, Clauw *et al* 2022). Moderately elevated temperatures that are below heat shock (27°C), as used in our experiments incorporating *myb59* mutants, do not trigger immediate stress responses, but may alter plant growth and development, e.g. root elongation, petiole hyponasty, and reduced stomatal index (Lee et al., 2021; Ludwig et al., 2021). Root elongation and transpiration rate increases in response to high temperatures regardless of watering conditions, highlighting the importance of a latent cooling capacity. Warm temperatures affect roots directly and indirectly, via decreases in shoot carbon availability and changes to root water relations due to increased transpirational demand, all of which can affect growth, water-, and nutrient uptake (Calleja-Cabrera et al., 2020). Root biomass and stomatal density have both been linked to temperature dependent acclimation of *A. thaliana* ecotypes. Ecotypes originating from sites with warmer temperatures allocated less biomass to the roots but showed a significant positive effect on reproductive allocation when grown under high temperatures (Vile et al., 2012). Here, local adaptations to warm environments appeared to be driven at the leaf level through reduced stomatal density, and via reduced biomass allocation to roots compared to ecotypes from colder sites when cultivated under high temperatures (Vile et al., 2012). On the other hand, high temperature environments have been shown to result in longer primary root length in exploration for water (Martins *et al*., 2017). We did not observe a significant difference in root length in wildtype plants grown under 22°C or 27°C, which may reflect the readily available water of plate grown plants (Supplemental Figure S10).

Given the context of the above-described trade-off between root elongation to facilitate water uptake for evaporative cooling and water conservation, the identification of *MYB59* via GWAS for VWU was especially interesting as this transcription factor is involved in cell cycle progression and root growth (Mu et al., 2009; Fasani et al., 2019). *MYB59* is expressed at higher levels in the roots specifically in xylem cells and might be involved in the transcriptional regulation of xylem differentiation (Oh et al., 2003). Over-expression of *MYB59* represses root growth (Mu et al., 2009, Enomoto et al 2023), and conversely, a *myb59* knockout mutant showed enhanced root elongation when grown on MS media containing 1% sucrose (Fasani et al., 2019). Reduced xylem differentiation in the roots has been shown to hinder water extraction altering drought responses (Tang et al. 2018), and root hydraulics plays a key role in maintaining the overall plant water status, especially in conditions of strong water demand (elevated temperatures) or water deprivation (Tang et al. 2018). Interestingly, we see a positive correlation between root hydraulic properties determined by Tang et al (2018) and integrated water use efficiency (δ^13^C) for the six ecotypes grown as part of our outdoor experiment (Supplemental Figure S3), highlighting the essential link between root water uptake and stomatal responses.

In our study, *myb59* mutants used significantly less water, and had lower water potential under the warm temperatures compared to wild type (Figure 6), which is likely due to a collation of factors including altered root growth, and reduced stomatal density, all of which may contribute to improved water homeostasis under elevated temperature (Supplemental Figures S7-11). The long root phenotype at elevated temperatures in *myb59* mutants is, however, counterintuitive to the reduction in VWU under elevated temperatures.

*MYB59* also plays a role in the balance of plant nutrient levels especially potassium (K^+^) and nitrate (NO_3_^-^). Under low K^+^/NO_3_^-^, the alternative splice variant MYB59α regulates the induction of the K^+^/NO_3_^−^ symporter NPF7.3, enhancing root-to-shoot translocation of K^+^ and over-expression of the *MYB59α* variant suppresses root growth under K^+^ replete conditions (Enomoto *et al*., 2023). Conversely, *myb59* mutants were shown to be defective in K^+^/NO_3_^-^ root-to-shoot translocation when grown on high levels of K^+^/NO_3_^-^, and under those conditions plants also exhibited an elongated root phenotype (Du et al 2019).

K^+^ movements are linked to water transport and accumulation of K^+^ in the vacuole allows plants to adjust water flux and cell water potential under water limiting conditions, and therefore a defect in K^+^ transport may limit water movements despite the long root phenotype. Furthermore, experiments to determine root length were carried out on plates under well-watered and high humidity conditions, while water use was determined in soil grown plants subjected to short-term dehydration and elevated temperatures. While root functional traits strongly influence water-use and nutrient uptake, plants may shift allocation among tissues to acquire resources that most limit growth depending on the environmental context, i.e., the need to conserve water (during short-term dehydration combined with elevated temperature), or the need for evaporative cooling (long term elevation of temperature).

## Conclusion

Our results suggests that *MYB59* regulates subtle root and stomatal adaptations of water use traits in connection with warmer environments (Figure 6, Supplemental Figures S7-11), but not necessarily short-term stress responses to elevated temperatures, and under cooler temperatures these differences were largely not significant. Functional *MYB59* appears linked to warm temperature adaptation due to its regulation of traits linked to water use and evaporative cooling and balancing nutrient and water requirements under prevailing environmental conditions. This is reflected in *MYB59* alleles demonstrating differential probability densities across temperature-associated bioclimatic parameters, such as mean annual temperature. This same effect was not observed with precipitation-associated parameters, highlighting the plausibility of this being a temperature adaptive mechanism (Figure 5).

## Materials and Methods

### Controlled growth environment experiment

78 *A. thaliana* ecotypes (Table S1) from the globally distributed Regional Mapping (RegMap) panel (Horton *et al*. 2012) were randomly selected and grown as part of the controlled growth environmental conditions experiments. Plants were grown in different temporal blocks over a period of 2 years where ecotypes Col-0 and C24 were common to each block. 10-15 biological replicates of each ecotype were grown. Plants were initially grown in a custom walk-in growth chamber before being transitioned to a glasshouse after the dry down experiment (see below), thereby replicating a short- to long-day transition. The conditions within both growing environments were exactly as previously described (Ferguson *et al*., 2018, 2019).

At 40 days post sowing, plants were subjected to a period of water withdrawal from ∼100% to ∼20% relative soil water content (rSWC) as described in Bechtold et al. 2016. The change in rSWC for each plant was modelled by a linear model to determine vegetative water use (VWU), (Ferguson *et al*., 2019).

Flowering time and the number of leaves at flowering was measured as the time point (day) of bud initiation. Upon opening of the final flower, plants were bagged and allowed to dry down. Above ground biomass was separated into three components: seeds, stalks and pod, and rosette. Harvest index was calculated as the ratio of seed biomass to total above ground biomass.

### Outdoor experiment

Six ecotypes (C24, Col-0, Ct-1, Est-1, HR-5, and Se-0) which were part of the original 78 ecotypes were selected based on their VWU profile for an outdoor experiment. After sowing, all seeds were stratified to promote germination. Initially, plants were grown in the above-described short-day controlled conditions as detailed previously (Ferguson *et al*., 2018, 2019). At 35 days post sowing, plants were transplanted into an outdoor planting bed (1m x 1m x 0.6m) filled with peat-based compost (Levington F2 + S, The Scotts Company, Ipswich, UK). The planting bed was located on the roof of the Life Science department at the University of Essex (51.87°N, 0.95°E). The outdoor experiment lasted 10 weeks from August 2015 until October 2015, where weather conditions did not deviate from average for this time of year (Supplemental Table S4). Plants were grown following a randomised design in rows and columns 8cm apart. 30 plants of each ecotype were grown.

Flowering and biomass accumulation were recorded at the end of the experiment as per the controlled environment experiments. In addition, we sampled a random fully expanded upper rosette leaf 5-days post bud initiation from five biological repeats of each ecotype per treatment. The sampling was ∼25 days post transplanting. Here, whole leaves were sampled immediately in liquid nitrogen to halt enzymatic activity and then used for measurements of leaf carbon content and carbon isotope composition (δ^13^C) via isotope ratio mass spectrometry as described previously (Roussel *et al*., 2009). δ^13^C was also determined on five biological repeats of the same ecotypes growing as part the above-described controlled growth conditions experiment.

### Statistical analysis and figure generation

All statistical analyses were performed within R. For all traits, we used the lme4 package to construct linear mixed models to explore sources of phenotypic variance where *Ecotype* and *Temporal block* were treated as random effects. From these, we estimated broad sense heritability (*H^2^*) as the ratio of the ecotype variation to the total observed variation. In addition, we extracted best linear unbiased predators (BLUPs) for each ecotype and added these to population means to generate predicted means (Supplemental Table S2) that controlled for the variation not due to ecotype.

We explored correlations between all traits via Pearson correlations, where p-values were Bonferroni corrected. We used the corrr package to visualize all pairwise correlations as a network, where traits were spaced according to multidimensional scaling. We further explored the relationship between the phenotypic traits in all ecotypes via principal component analysis (PCA) using the FactoMineR package, where data was scaled to unit variance. Scree plots were inspected to determine significant principal components (PCs), which were defined as those that explained more variance than would be expected by chance (Kaiser, 1991). To identify representative ecotypes, we first computed a distance matrix using Euclidean measures based on the entire phenotypic data set. We then used the Ward2 agglomeration to perform a hierarchical clustering analysis. The hierarchical clustering was visualized using a fan dendrogram where five groups were chosen for cutting the tree.

Climatic data from the point of origin of all ecotypes were extracted using the R package raster. This generated a dataset consisting of 18 biologically relevant climatic (BIOCLIM) variables based on latitude and longitude (Hijmans, 2012). PCA was performed on the climatic dataset as described for the trait dataset. Following the methodology of (Wolfe & Tonsor, 2014), we explored then correlations between significant trait PCs (TPCs) and climatic PCs (CPCs) as evidence of climatic selection on phenotypic variance. This was achieved by performing Pearson’s product moment correlation testing between each TPC-CPC pairwise interaction.

Simple linear statistical models were constructed to test for interactions between traits that were measured in common between the controlled environment experiment and the outdoor experiment.

All figures were produced within R using base plotting functions and ggplot2. Additional post-processing was performed in Affinity Designer.

### GWAS and population genetics analysis

We used the R package adegenet (Jombart, 2008) to group ecotypes based on genetic similarity using the 250k RegMap single nucleotide polymorphism (SNP) dataset (Horton *et al*., 2012). Initially, the find.clusters() function was used to transform the SNP matrix via PCA before running iterative K-means clustering. The most appropriate clustering model was selected according to the Bayesian information criterion (BIC). Subsequently, the dapc() function was used to perform discriminant analysis of principal components (DPAC; (Jombart *et al*., 2010)) to define the diversity between the pre-identified groups. We performed Kruskal-Wallis rank sum tests to test for significant differences between in mean annual temperature and VWU between the identified DAPC groups.

Genome-wide association mapping (GWAS) was performed for VWU with GWAPP (Seren *et al*., 2012), which uses ∼206,000 SNPs generated from ecotypes that were genotyped for 214,000 SNPs and/or sequenced via next generation sequencing. We acknowledge that incorporating <100 populations into GWAS analyses is typically cautioned against due to reduced statistical power. However, given instances where population number is restricted it is still possible to associate genetic and phenotypic variation, especially when trait variation is substantial (Lehnert *et al*., 2018; Soumya *et al*., 2021)To this end we note the success of other Arabidopsis studies that have incorporated low numbers of populations (70-150, Zou et al., 2017; de Ollas et al., 2023; Wang et al., 2024). In addition, we have carried out GWAS according to best practices, used a stringent significance criterion for identifying trait associated SNPs and performed validation on a candidate gene. The accelerated mixed model (AMM) genome scan approach (Zhang *et al*., 2010) was used for our GWAS. We corrected the p-values according to (Benjamini & Hochberg, 1995) to identify SNPs below a 5% false discovery threshold. The linkage disequilibrium (LD) block (*r^2^*> 0.3) containing the significant SNPs was investigated to identify candidate genes using GWAPP.

AT5G59780 (*MYB59*) was identified as a candidate gene for VWU based on GWAS results. To validate the effect of *MYB59* on VWU, we generated haplotypes of *MYB59* based on the seven SNPs from the 250K RegMap dataset (Horton *et al*., 2012; Supplemental Table 6) that fall within the coordinates of *MYB59*. Significant differences in representative trait and BIOCLIM parameters between the two main haplotype groups were tested by fitting a one-way ANOVA.

To test how natural allelic variation in *MYB59* was associated with temperature variation we employed the approach of (Ferrero-Serrano & Assmann, 2019) to test for associations between genetic variants with local environment. Here, we used the Polymorph variant browser (www.1001genome.org/tools) to search for SNPs with a minor allele frequency >5% within *MYB59* with a predicted “moderate” or greater impact on the translated amino acid sequence. Subsequently, we downloaded the *MYB59* sequence data for all ecotypes from the 1001 genome initiative and filtered for those pre identified SNPs. These SNPs were visualised on the *MYB59* gene model using the R package genemodel (https://github.com/greymonroe/genemodel). We tested the probability density of each allele of these SNPs against variation in the BIOCLIM parameters for all ecotypes comprising the 1001 genome initiative. We omitted sequences of ecotypes from North America and the British Isles to avoid confounding effects due to recent dispersal that have not allowed enough time for selection to have occurred (Platt *et al*., 2010). Observed differences were tested for significance via two-sample Wilcoxon tests.

### *myb59* knockout experiments

Based on the GWAS results, AT5G9780 (*MYB59*) was selected for experimental investigation. Two previously published myb59 knockouts (*myb59-1*, *myb59-2*; Fasani et al., 2019; Du et al., 2019) were obtained from Drs Wang and Furini. Mutants were checked for the presence of the T-DNA insertion (myb59-1) and MYB59 expression using RT-PCR (Primers: myb _fw2: GGGCGGTCTTTGGAAAGGACC and myb_rev2 CTAAAGGCGACCACTACCATG, Supplementary Figure x). The *myb59-2* allele was generated using CRISPR/Cas9, and maintains a full-length transcript, however phenotypically the mutant copies the *myb59-1* mutant, suggesting that the protein is not functional (Du et al 2019).

Seeds from both knockouts and Col-0 wildtype were sown as described above and short dehydration experiments were carried out at 22⁰C (day/night) and 27⁰C/22⁰C (day/night) short-day conditions to determine VWU. Water potential was measured using a WP4C Water Potential Meter (Labcell, Ltd, UK) on leaf disks from 6 independent plants at the mid-point of the light period. For stomatal densities, the abaxial side of the leaf was painted with clear nail varnish and the peels were analysed for stomatal densities using a Zeiss Axioskop 20x magnification connected to a Retiga 2000R camera. Additionally, to the above-described experiment, Col-0, *myb59-1* and *myb59-2* alleles mutants were sown on ½ MS media to assess root length at 22⁰C (day/night) and 27⁰C/22⁰C (day/night) conditions. Roots were measured using ImageJ at every 2 days post germination up until 2 weeks after sowing. Leaf temperature was measured using an infrared thermal imaging camera (HIKMIKRO) before the start of the VWU measurements.

## Contributions

UB and JNF designed and performed the experiments and analysed the data. JNF and UB wrote the manuscript with input from all authors. OB performed the carbon and δ^13^C data analysis.

## Acknowledgements

The authors thank Susan Corbett and Philip Davey for help with general plant maintenance and help with setting up the outdoor experiment. JNF was supported by a BBSRC-CASE studentship (BB/J012564/1). The authors thank Prof Furini and Prof Wang for providing seeds of the *myb59-1 and myb59-2* knockout lines, respectively. The authors would like to thank SILVATECH (Silvatech, INRAE, 2018. Structural and functional analysis of tree and wood Facility, doi: 10.15454/1.5572400113627854E12) from UMR 1434 SILVA, 1136 IAM, 1138 BEF and 4370 EA LERMAB from the research centre INRAE Grand-Est Nancy for the carbon stable isotope analysis. SILVATECH facility is supported by the French National Research Agency through the Laboratory of Excellence ARBRE (ANR-11-LABX-0002-01).

## Data availability

All data are available in the supporting information and on request to the corresponding author.

## Supplemental Material

Supplemental Figure S1. Comparison of *R^2^* between linear models and segmented models

Supplemental Figure S2. Variation of all traits measured as part of controlled environment experiment

Supplemental Figure S3. Comparison on integrated water-use efficiency (δ^13^C) measured in this study and leaf hydraulics (*L*p_r_) measured in Tang *et al*. (2018)

Supplemental Figure S4. Scree plots for (A) TPCA and (B) CPCA

Supplemental Figure S5. Linkage disequilibrium block containing AT5G59780 (*MYB59*) and the two significant SNPs from GWAS for VWU

Supplemental Figure S6. MYB59 haplotype analysis

Supplemental Figure S7. Rosette areas of wildtype and *myb59* knockout lines

Supplemental Figure S8. Stomatal density of wildtype and *myb59* knockout lines

Supplemental Figure S9. Root length of wildtype and *myb59* knockout lines 12 days post sowing

Supplemental Figure S10. (A) Root biomass and (B) root:shoot ratio of wildtype and *myb59* knockout lines grown at 27⁰C

Supplemental Table S1. Ecotypes comprising study

Supplemental Table S2. Predicted means for all traits

Supplemental Table S3. Correlations of traits measured as part of the controlled environment experiments

Supplemental Table S4. Comparisons of climatic parameters and phenotypic traits between DPAC groups

Supplemental Table S5. Results from GWAS for VWU

Supplemental Table S6: SNPs from the AMM GWAS. Significant SNPs are highlighted in orange

Supplemental Table S7. Trait and climatic parameters of MYB59 haplotype groups

