## Supplementary figures and images for "MYB59 is linked to natural variation of water use associated with warmer temperatures in *Arabidopsis thaliana*"

### Figure S1

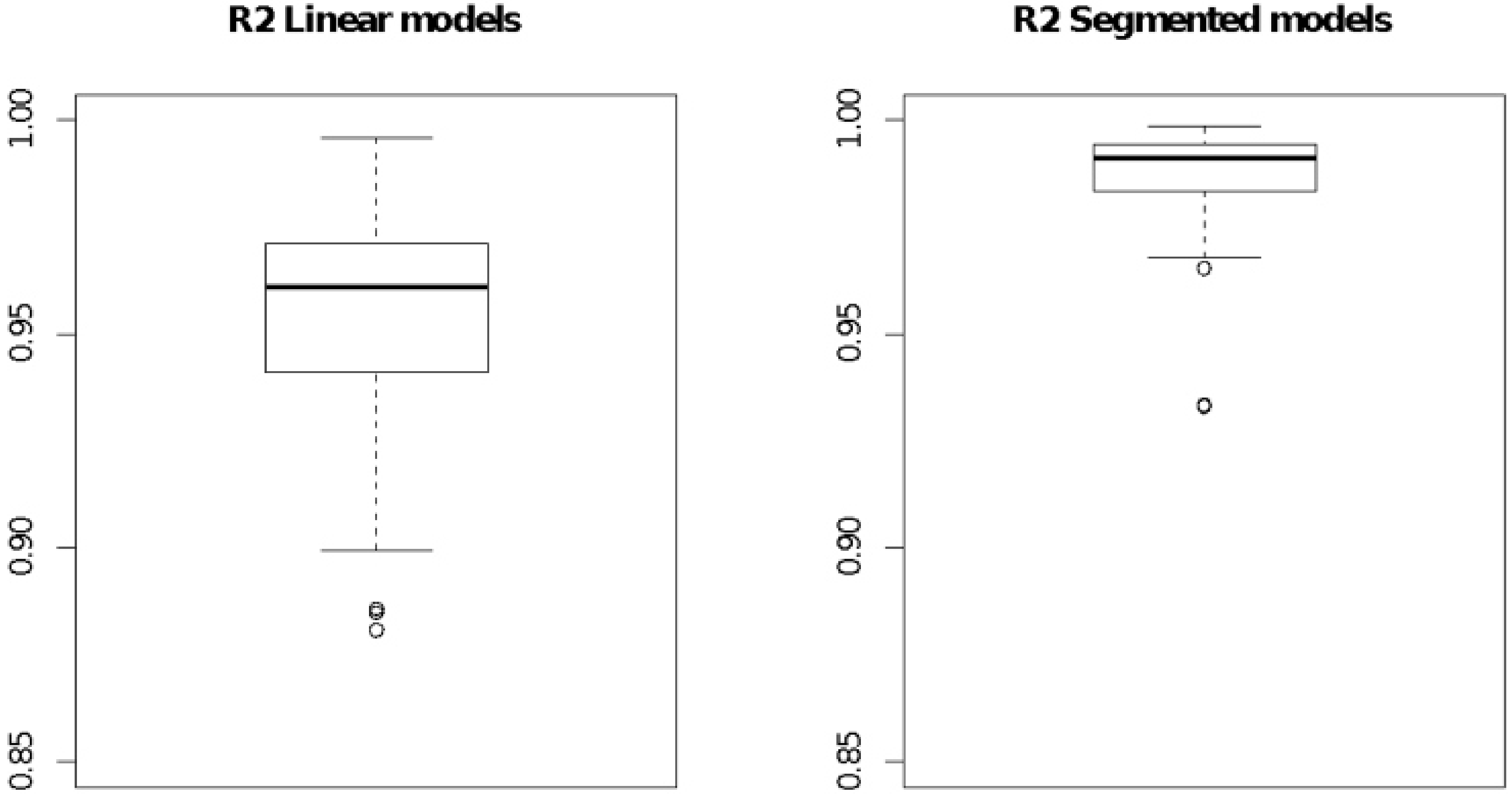


Figure S1 Ferguson et al

### Figure S2

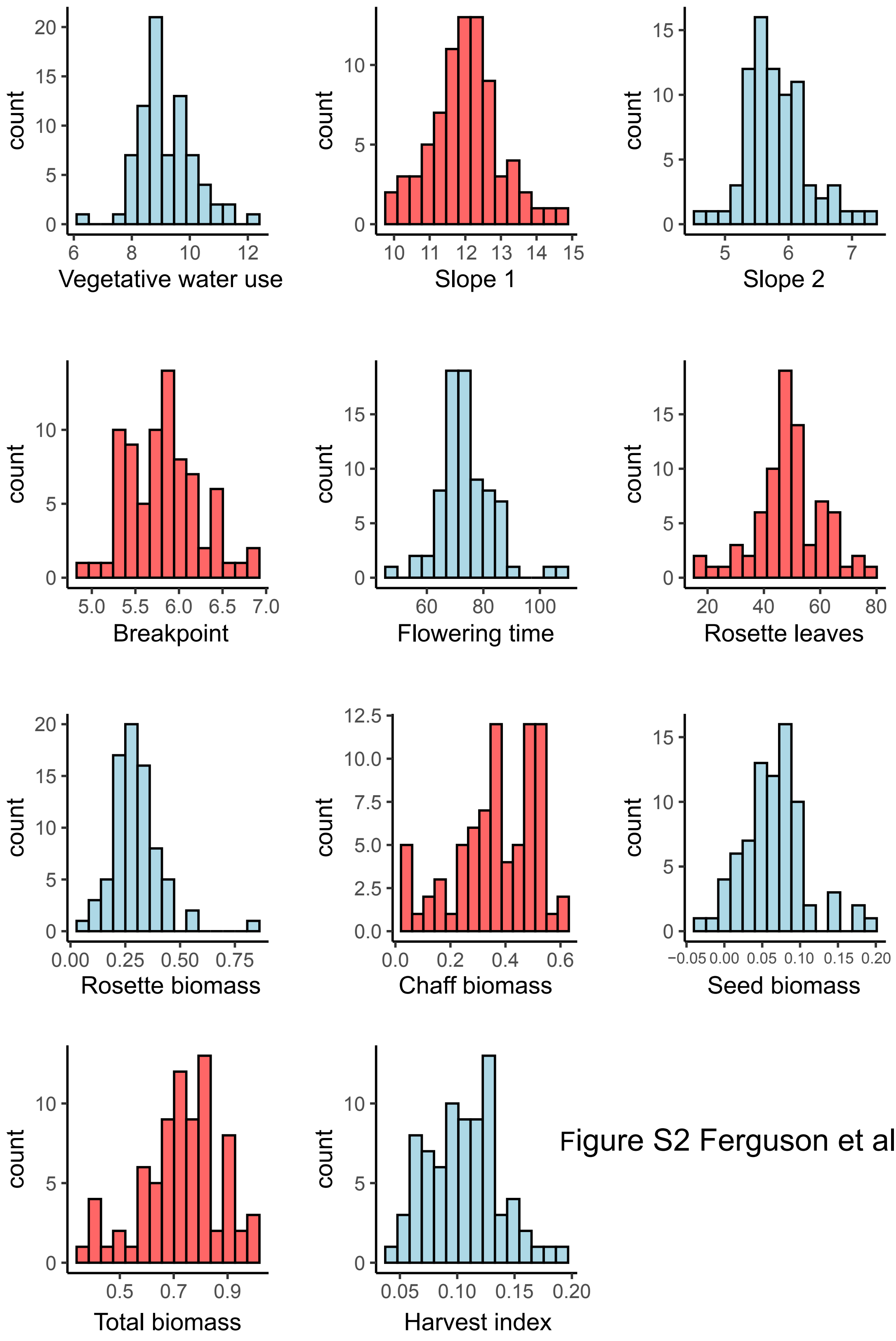


Figure S2 Ferguson et al

### Figure S3

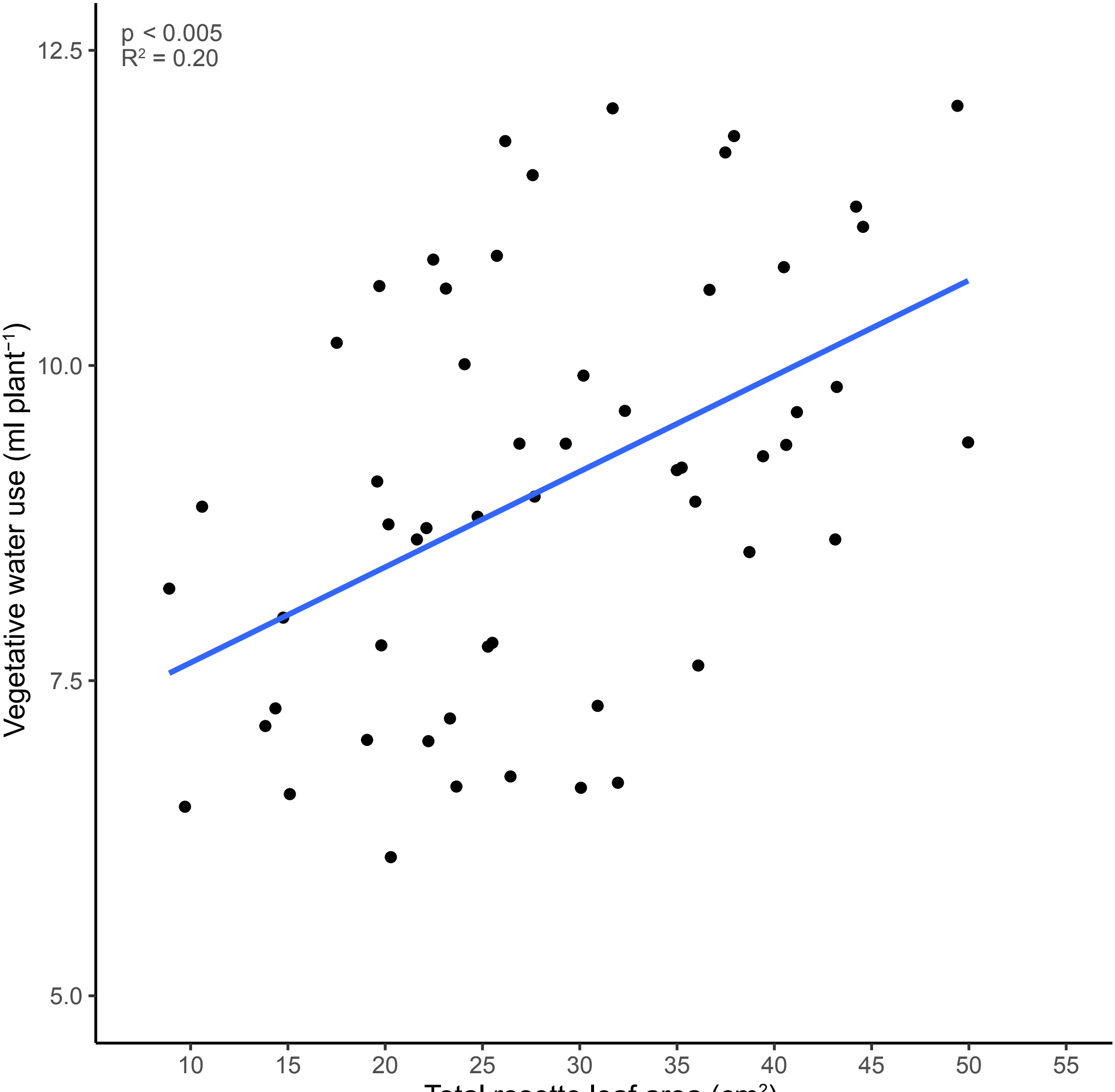


Figure S3 Ferguson et al

### Figure S6

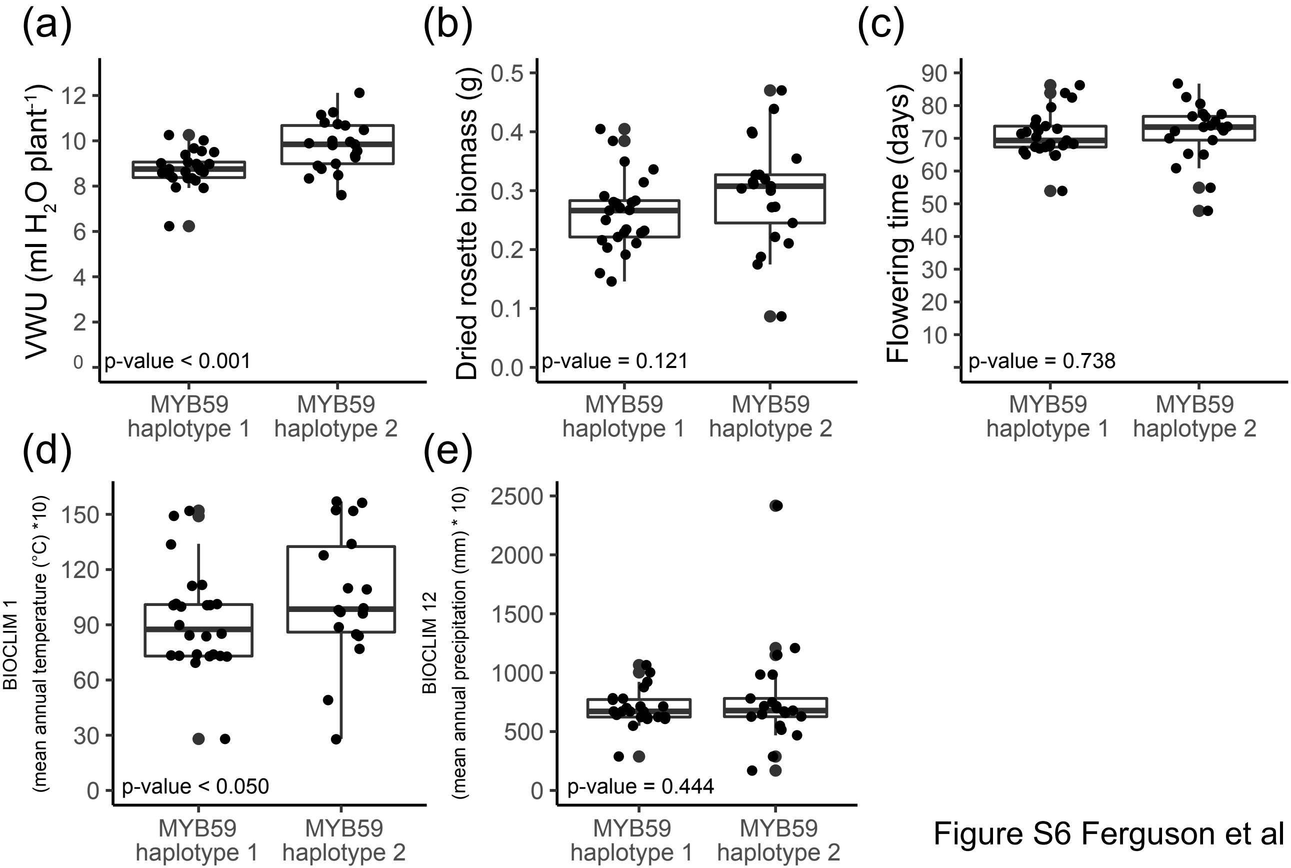


Figure S6 Ferguson et al

### Figure S7

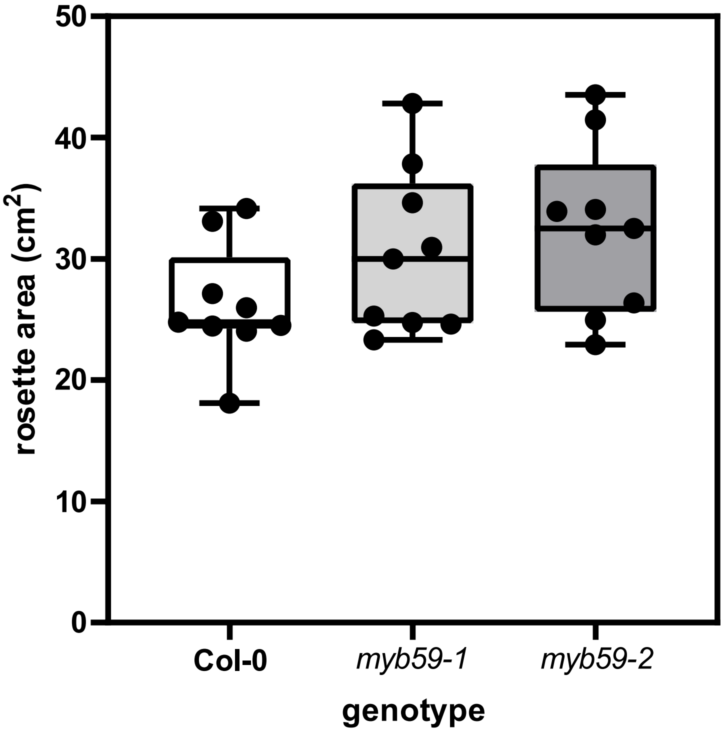


Figure S7 Ferguson et al

### Figure S8

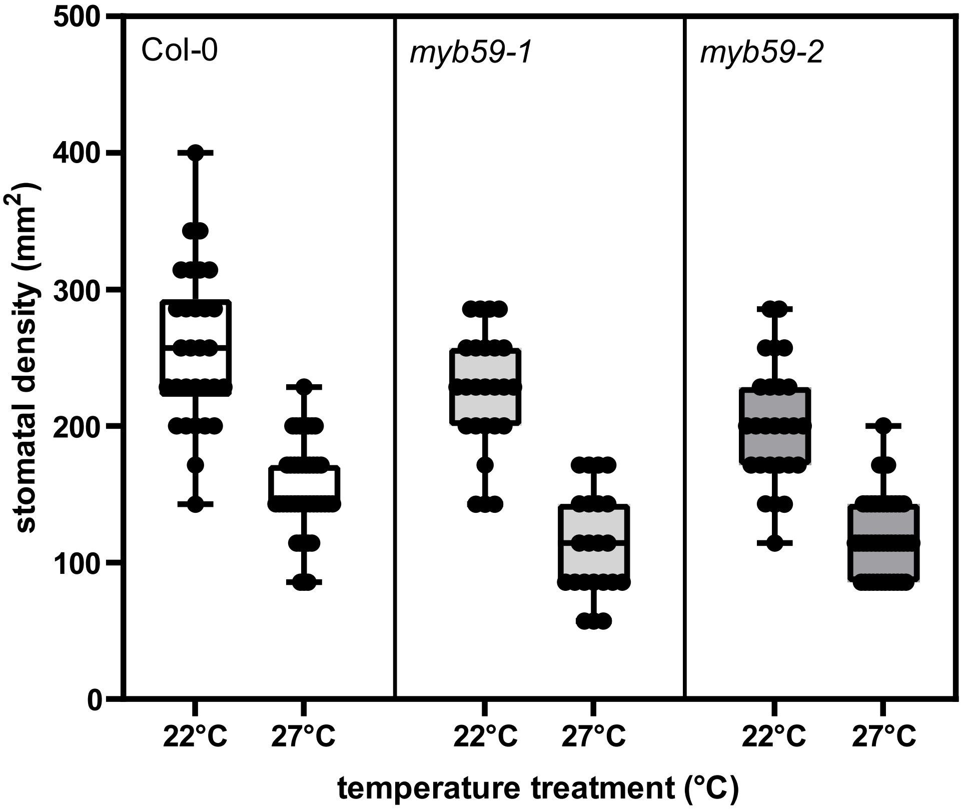


Figure S8 Ferguson et al

### Figure S10

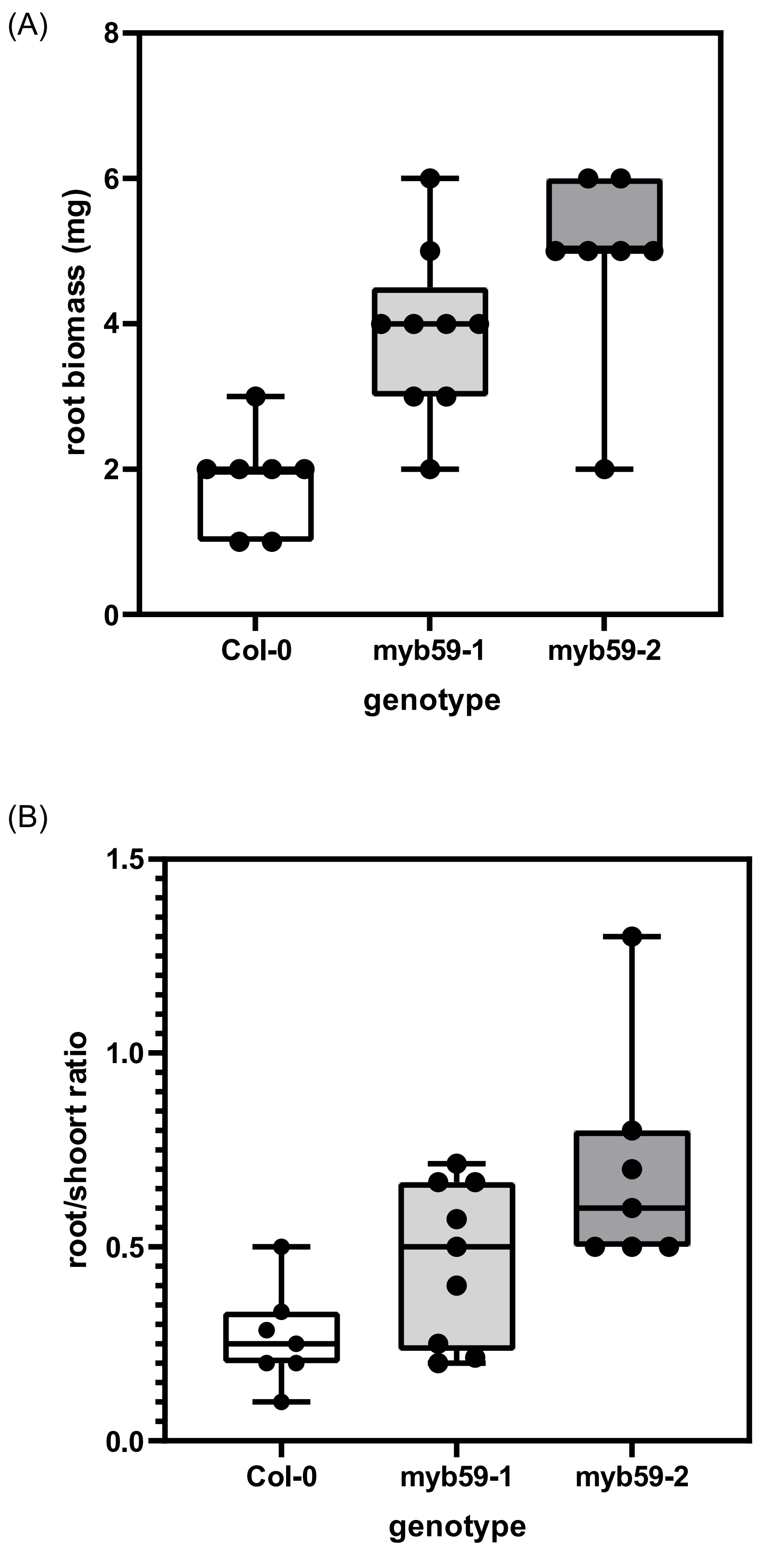


Figure S10 Ferguson et al
