## Supplementary material for "MYB59 is linked to natural variation of water use associated with warmer temperatures in *Arabidopsis thaliana*": Figure S5

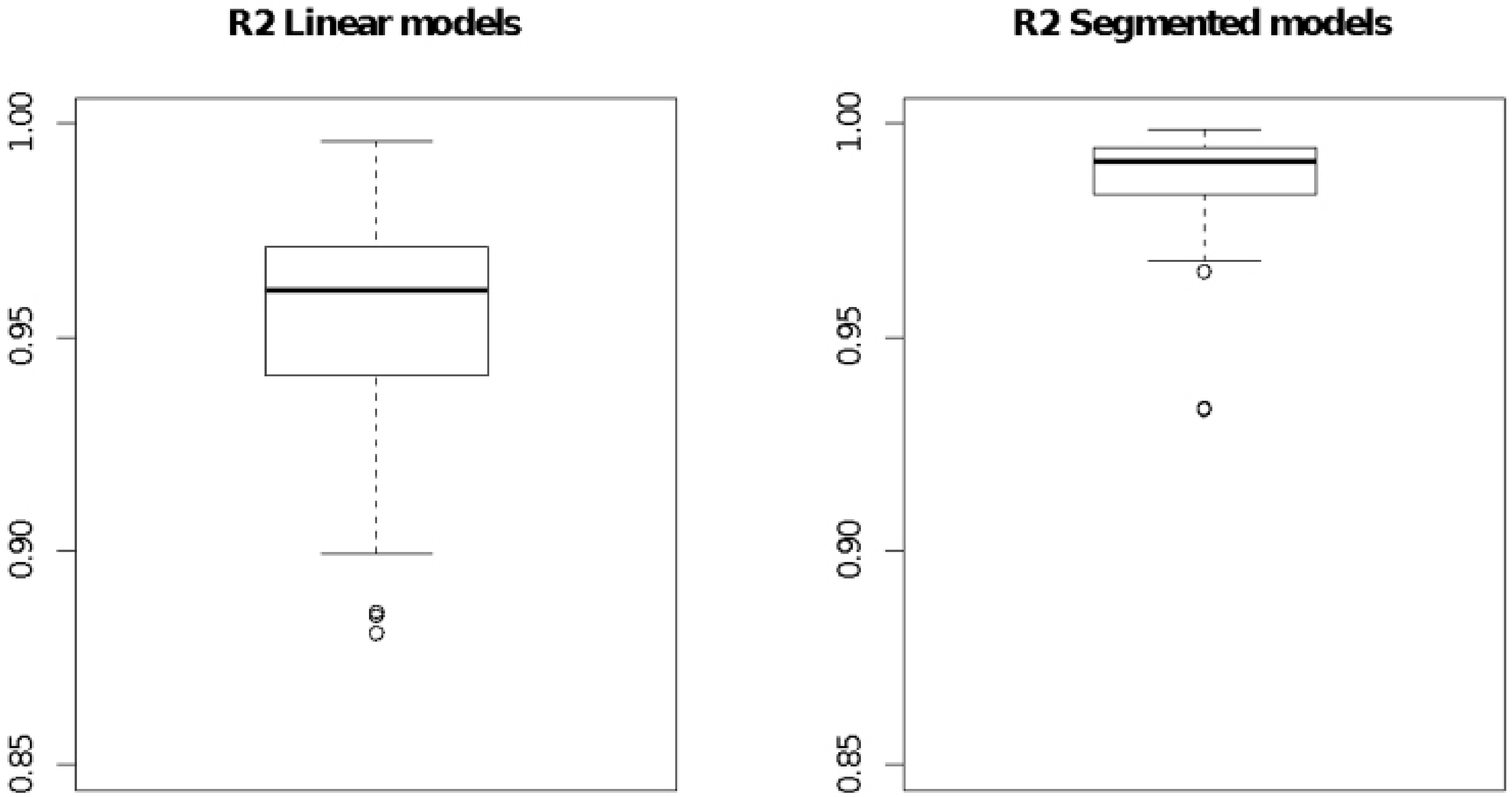


Figure S1 Ferguson et al

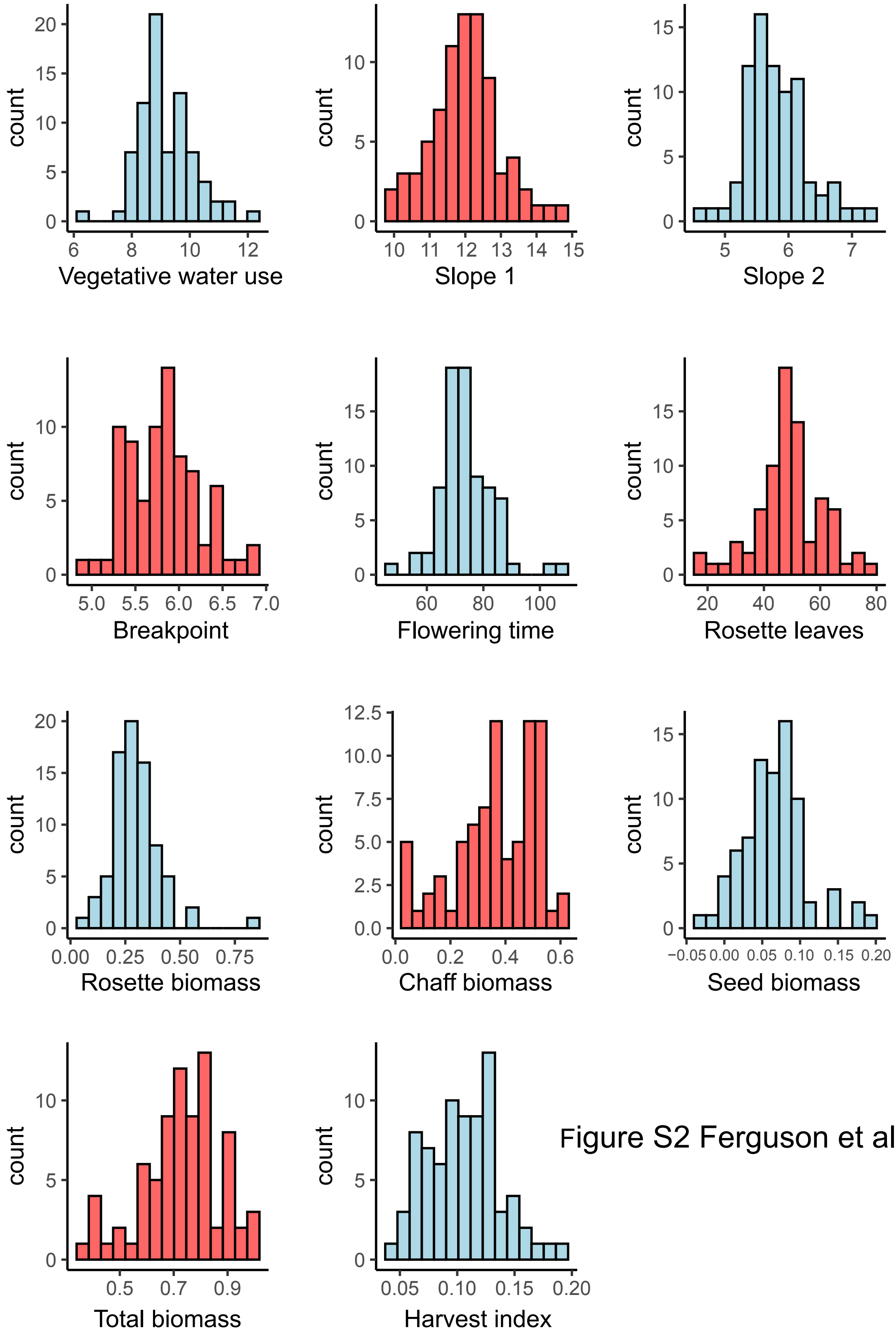


Figure S2 Ferguson et al

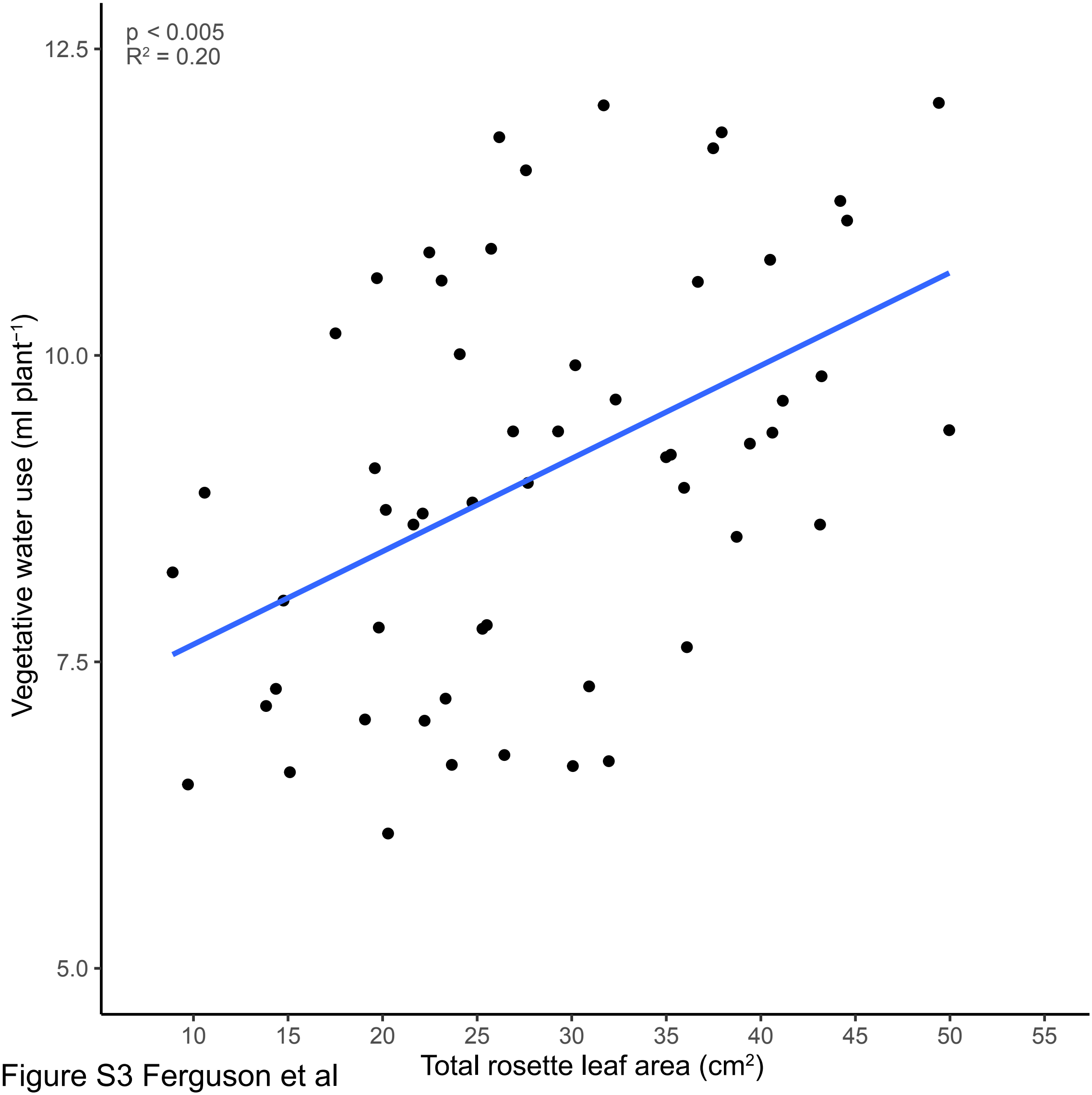


Figure S3 Ferguson et al

Figure S4 Ferguson et al

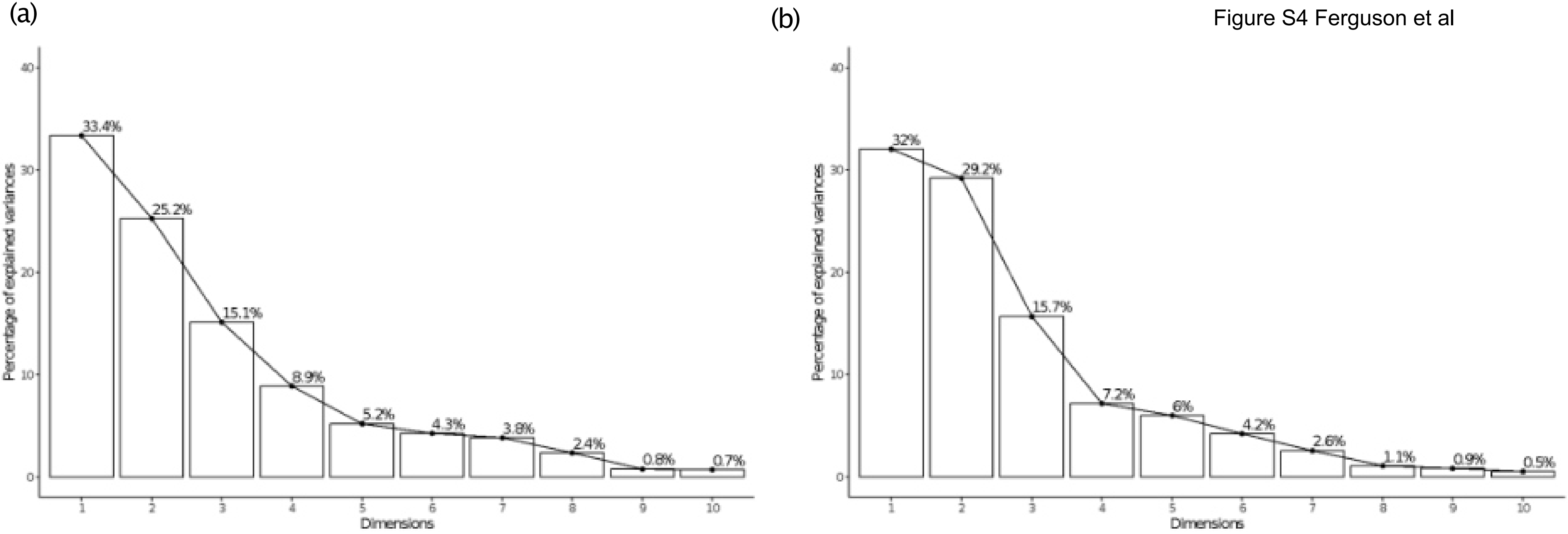

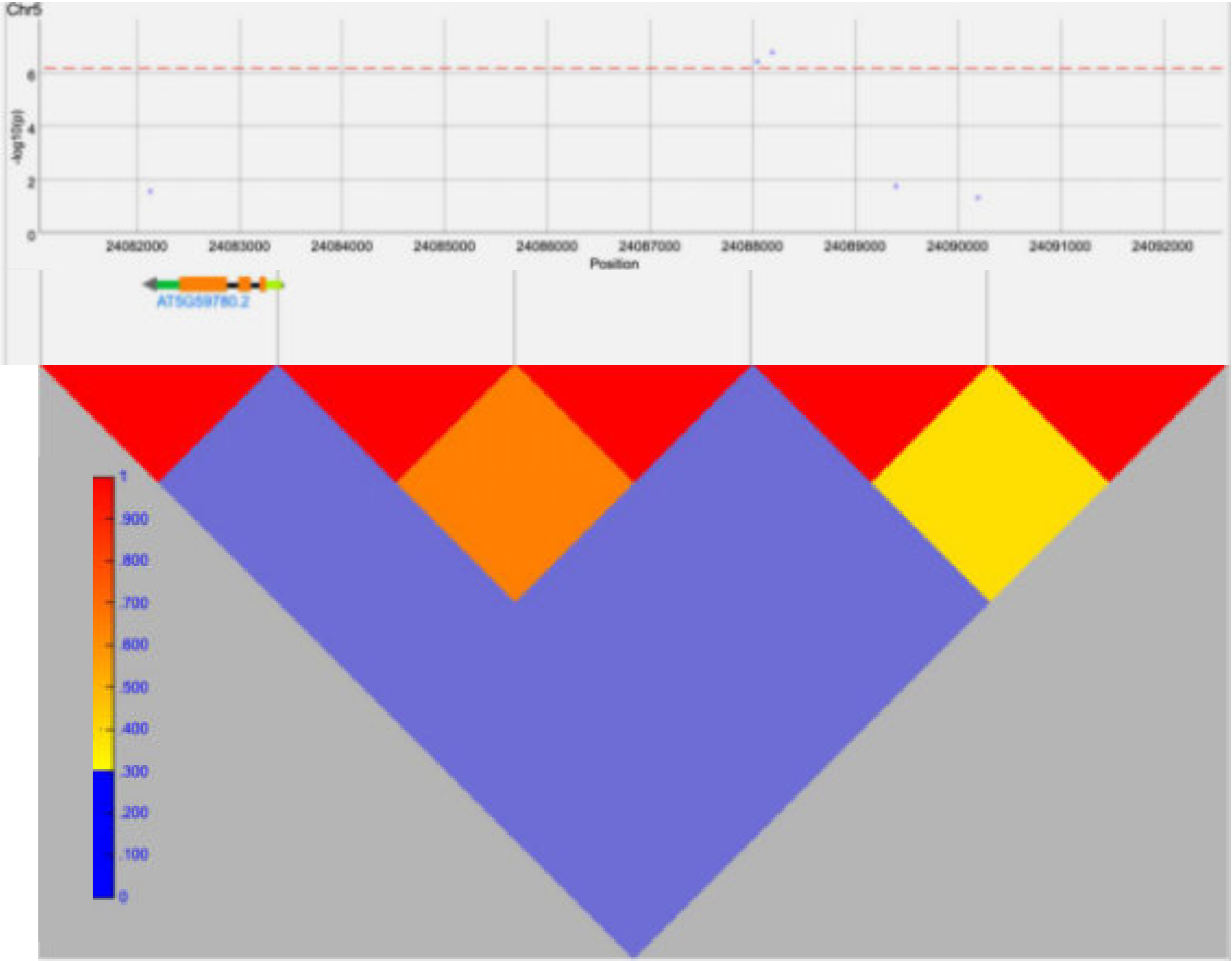


Figure S5 Ferguson et al

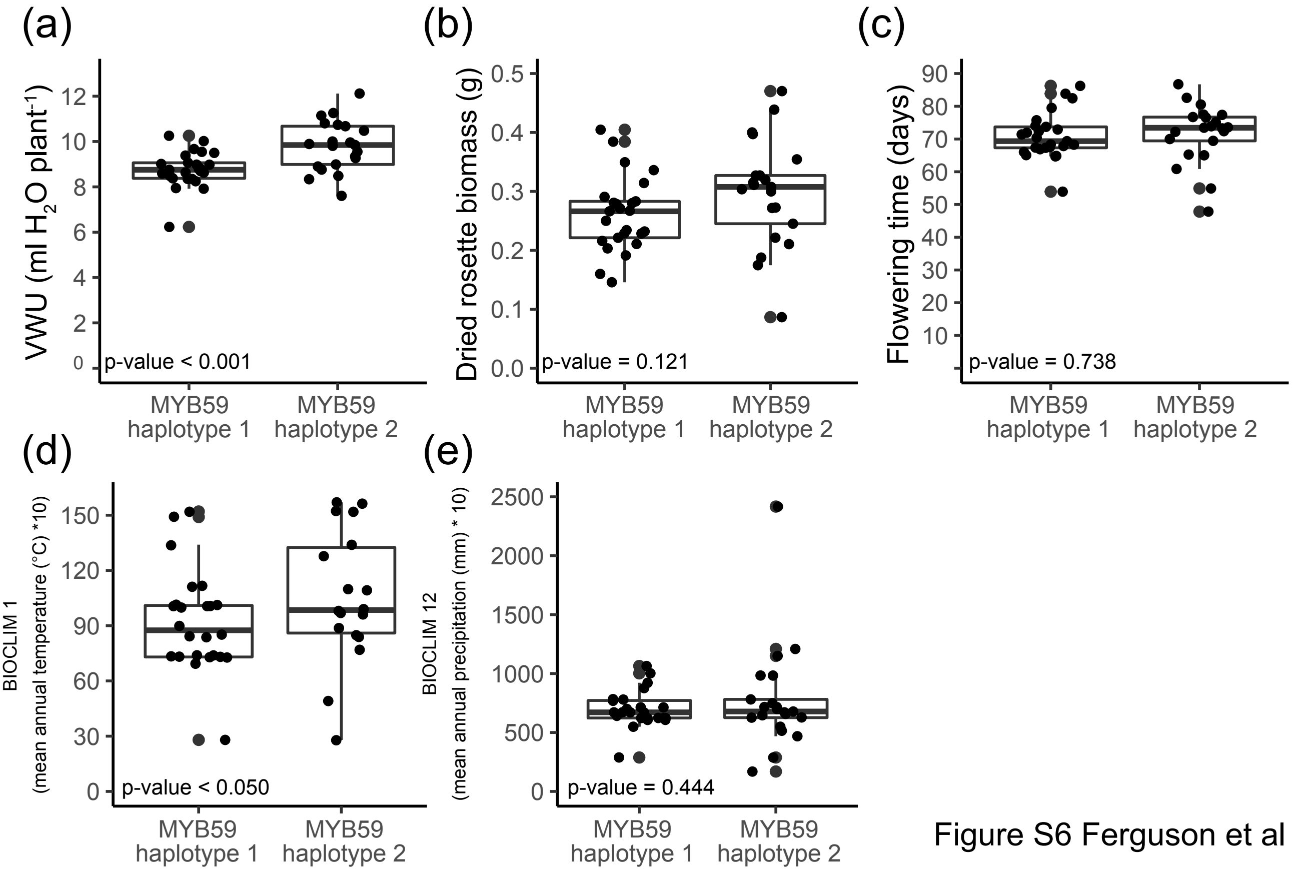


Figure S6 Ferguson et al

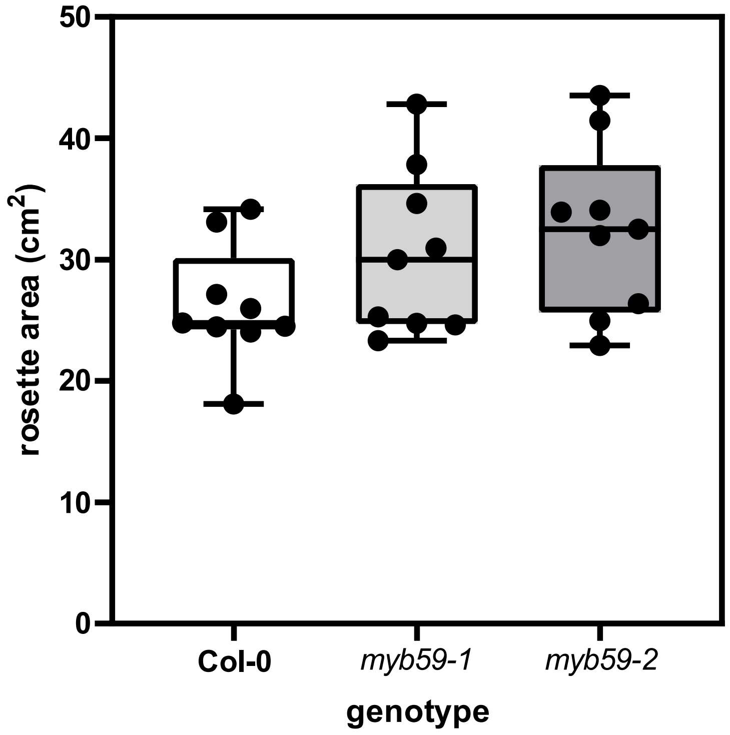


Figure S7 Ferguson et al

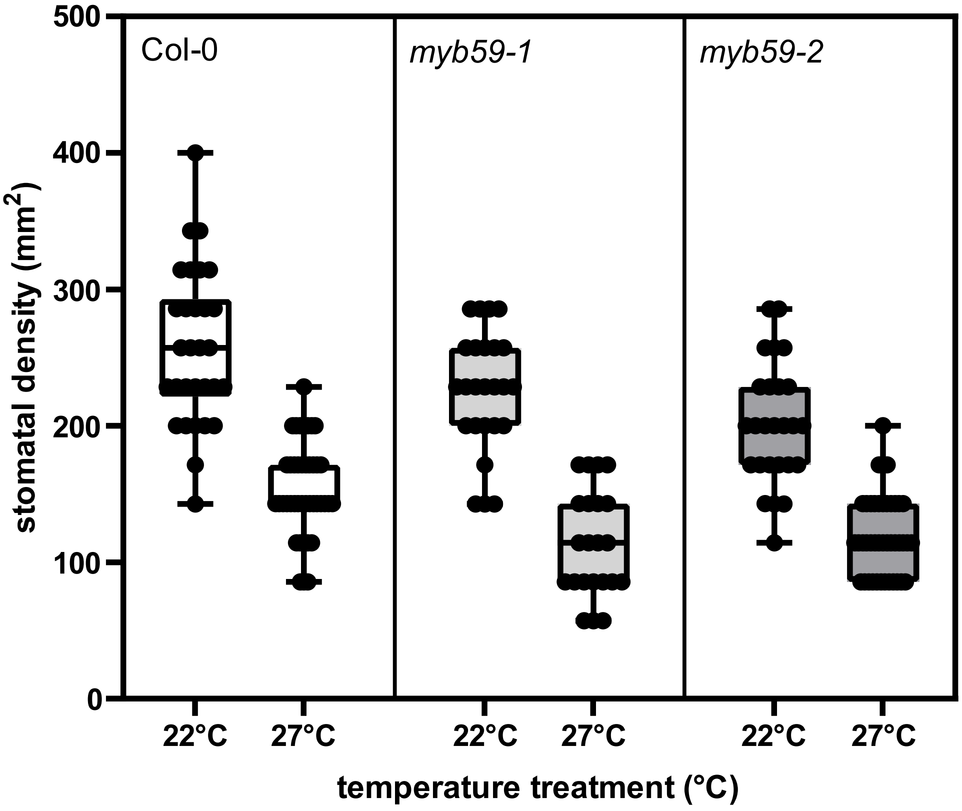


Figure S8 Ferguson et al

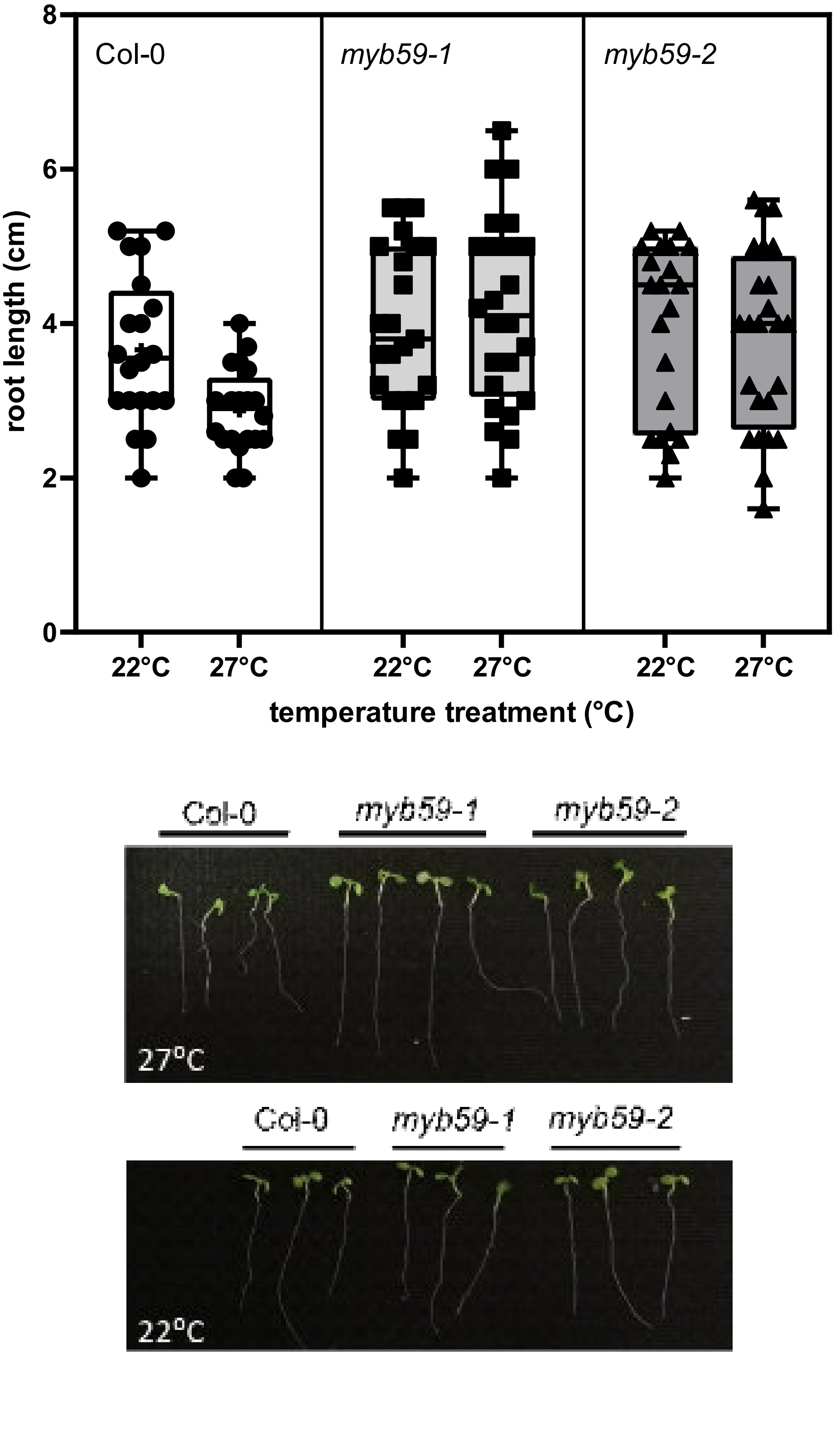


Figure S9 Ferguson et al

(A)

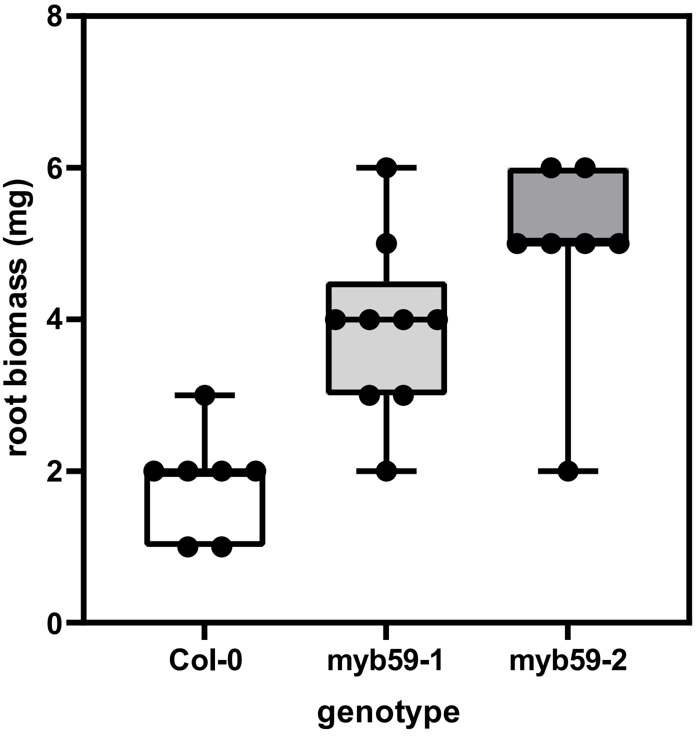


(B)

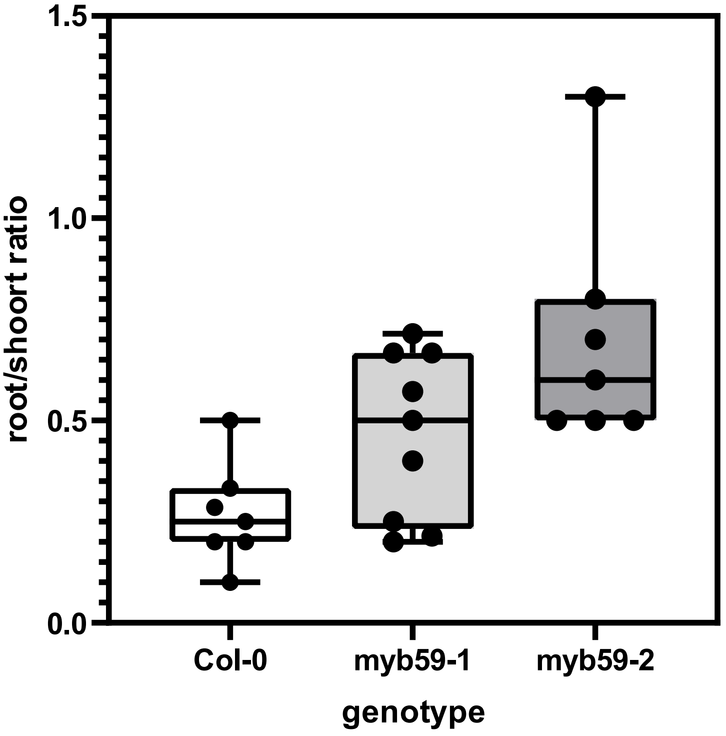


Figure S10 Ferguson et al
